# Spindle - slow oscillation coupling correlates with memory performance and connectivity changes in a hippocampal network after sleep

**DOI:** 10.1101/2021.10.27.466071

**Authors:** Lisa Bastian, Anumita Samanta, Demetrius Ribeiro de Paula, Frederik Weber, Robby Schoenfeld, Martin Dresler, Lisa Genzel

**Affiliations:** Donders Institute for Brain Cognition and Behaviour Radboud University, Postbus 9010, 6500GL Nijmegen, Netherlands; Donders Institute for Brain Cognition and Behaviour Radboud University Medical Centre; University of Halle

**Keywords:** memory consolidation, resting-state networks, sleep, spindles, slow oscillations

## Abstract

After experiences are encoded, post-encoding reactivations during sleep have been proposed to mediate long-term memory consolidation. Spindle-slow oscillation coupling during NREM sleep is a candidate mechanism through which a hippocampal-cortical dialogue may strengthen a newly formed memory engram. Here, we investigated the role of fast spindle- and slow spindle-slow oscillation coupling in the consolidation of spatial memory in humans with a virtual water maze task involving allocentric and egocentric learning strategies. Furthermore, we analyzed how resting-state functional connectivity evolved across learning, consolidation, and retrieval of this task using a data-driven approach. Our results show task-related connectivity changes in the executive control network, the default mode network, and the hippocampal network at post-task rest. The hippocampal network could further be divided into two subnetworks of which only one showed modulation by sleep. Decreased functional connectivity in this subnetwork was associated with higher spindle-slow oscillation coupling power, which was also related to better memory performance at test. Overall, this study contributes to a more holistic understanding of the functional resting-state networks and the mechanisms during sleep associated to spatial memory consolidation.

## Introduction

Systems consolidation theory is one of the most widely studied frameworks of long-term memory formation. The theory posits that labile engrams in the brain, encoded during learning, are strengthened through repeated reactivation. Over time, this process might lead to a more stable long-term representation – a consolidated memory trace (Squire et al., 2015). Reactivations of the engram are thought to occur predominantly during sleep but can also be seen during quiet rest. A potential key mechanism underlying systems consolidation constitutes the hierarchical coordination of three event types during non-rapid eye movement (NREM) sleep: (1) high-amplitude cortical slow oscillations, (2) thalamically-driven spindles, and (3) hippocampal sharp wave-ripples. The orchestrating of these oscillations driven by the waxing and waning of slow-oscillations in the prefrontal cortex (Helfrich et al., 2019) is thought to set ideal time windows for hippocampal information transfer to the cortex (Sirota et al., 2003) and brain-wide synaptic consolidation processes underlying systems consolidation (Genzel, 2020; Genzel, Kroes, et al., 2014; Navarro-Lobato & Genzel, 2020).

In humans, two kinds of spindles have consistently been identified: slow spindles (∼ 9-12.5 Hz) and fast spindles (∼ 12.5-16 Hz), which are most prominent over frontal and centro-parietal regions, respectively (Adamczyk et al., 2015; Anderer et al., 2001). Both spindle types are coupled to slow oscillations. While fast spindles are often nested in the slow oscillation’s excitable up-state, slow spindles typically occur during the transition period into the slow-oscillation’s inhibitory down-state (Molle et al., 2011). Fast spindles have been implicated as a facilitator of information transfer between hippocampus and neocortex as they coincide with hippocampal sharp wave-ripples (Staresina et al., 2015). Slow spindles, which are mostly preceded by ripples, may be related to a cortical cross-linking of transferred information and the modulation of prefrontal circuitry to form a neocortical memory representation (Molle et al., 2011). However, since only in human studies there is a consistent split between slow and fast spindles, while in studies with mice and rats this subdivision is rarely done, concrete evidence for a different function of each spindle type is lacking.

The influence of these two spindle types on behavior is even less conclusive as not many studies differentiate between slow and fast spindles. Recent EEG findings have shown that a larger amount and increased consistency in the coupling of fast spindles and slow oscillations is related to increased declarative memory performance (Hahn et al., 2020; Muehlroth et al., 2019; Niknazar et al., 2015). Experimental reinforcement of the coordination between hippocampal sharp wave-ripples, slow oscillations, and frontal spindles in rodents – potentially the analog of human slow spindles – also resulted in increased memory performance (Maingret et al., 2016). In contrast, preliminary findings on slow spindle-slow oscillation coupling in humans point towards decreased declarative memory performance with increased slow spindle-slow oscillation coupling or no effect at all (Barakat et al., 2011; Muehlroth et al., 2019). Thus, it is crucial to disentangle the behavioral relevance of both spindle types in memory consolidation.

Given the importance of sleep and the hippocampus in offline consolidation, it has been hypothesized that sleep plays a distinct role in the consolidation of hippocampus-dependent in contrast to ‘non-hippocampus-dependent’ memories. These two types of memories can be investigated using spatial navigation paradigms. Navigation in an environment involves the use of different navigational strategies, namely allocentric and egocentric strategies (for a review Ekstrom et al., 2017). In an allocentric reference frame, the navigator can infer their position based on the relative positions of stable landmarks (Klatzky, 1998). Allocentric spatial learning is suggested to strongly rely on the hippocampus as it requires a spatial cognitive map of the landmark positions (O’Keefe & Nadel, 1978). In contrast, the egocentric reference frame continuously changes with the movement of the navigator. During egocentric spatial learning, the navigator needs to keep track of the moved distance and the direction for which the dorsal striatum has been identified as a critical structure (McDonald & White, 1994). It may thus be proposed that, in a spatial context, consolidation mechanisms differ between allocentric and egocentric training conditions and influence memory performance accordingly. However, recent evidence points toward hippocampal involvement in consolidation during sleep, even for previously considered non-hippocampus-dependent memories (Sawangjit et al., 2018; Schapiro et al., 2019), and similar functional network changes after sleep across allocentric and egocentric training (Samanta et al., 2021). Therefore, it remains an open question how spatial information of the two reference frames is consolidated during sleep and how these mechanisms influence memory performance.

Although much work has focused on the role of sleep in systems memory consolidation, studies in both rodents (Davidson et al., 2009; Diba & Buzsáki, 2007) and humans (Schuck & Niv, 2019; Tambini & Davachi, 2013) have demonstrated system-wide consolidation processes during awake resting periods immediately after learning. Similar to sleep-dependent consolidation, event representations are spontaneously reactivated in the hippocampal-thalamo-cortical network during post-encoding rest (for a detailed review, see Tambini & Davachi, 2019). Important nodes of these reactivations are thought to incorporate the hippocampal network centered on the medial temporal lobe, medial temporal cortex, and retrosplenial cortex (van Buuren et al., 2019), as well as interactions between hubs of the default mode network (Lin et al., 2017) and task-relevant networks (e.g., the executive control network Sneve et al., 2017). Greater levels of resting-state functional connectivity between areas specific to the stimuli and the hippocampus were shown to predict the future ability to retrieve a memory (Tambini et al., 2010). In addition, functional connectivity immediately after learning as well as the amount of hippocampal reactivation predicted subsequent overnight memory retention, suggesting a complementary relationship between awake resting reactivation and later consolidation during sleep (Schapiro et al., 2018). While most previous studies separately examine one type of post-encoding period (i.e., rest or sleep), studies investigating consolidation processes across both periods are necessary to gain a holistic understanding of the parallels between mechanisms of consolidation during these distinct brain states.

The present study thus aims to provide within-study measures at different post-encoding brain states and show their relationship with memory performance in a spatial navigation task. Our study follows up on our investigations into the effect of sleep on allocentric and egocentric memory representations in humans and rats (Samanta et al., 2021). Participants performed a human analog of the water maze (Morris, 1981), a validated paradigm to study different aspects of spatial navigation (Muller et al., 2018; Schoenfeld et al., 2017). Whereas Samanta et al. (2021) focused on task-fMRI data during learning and retrieval, here we focused on resting-state fMRI and sleep-EEG data. Thus, resting-state functional connectivity in relevant networks was assessed before and after learning and retrieval of the memory task in which participants either engaged in allocentric or egocentric spatial navigation. EEG was used between the two task sessions to measure NREM sleep-related consolidation processes (i.e., spindle and slow oscillation properties and their coupling). We adopted a data-driven approach (Ribeiro de Paula et al., 2017) to investigate the evolution of resting-state functional connectivity across spatial learning, consolidation, and memory retrieval. Resting-state networks were selected based on the results obtained by Samanta et al. (2021), including the default mode network, the hippocampal network, and the right executive control network. To establish a triad relationship between sleep, resting-state and behavior, we correlated spindle and slow oscillation parameters, changes in resting-state functional connectivity, and performance measures of the spatial memory task.

Samanta et al. (2021) found similar sleep-related brain-wide changes at memory retrieval, for allocentric and egocentric training. Specifically, activations increased in the executive control network and decreased in the default mode network from learning to retrieval, if subjects slept after learning. Hence, we expected to observe similar changes in these two networks at rest. Brain-wide consolidation processes should also be observable for the hippocampal network that coordinates consolidation during sleep and at rest. We hypothesized that spindle-slow oscillation coupling is positively associated with changes in the hippocampal network and memory performance. These findings should not differ between memory conditions since Samanta et al. (2021) did not observe a difference between allocentric and egocentric training. Indeed, our results demonstrate that the executive control and the default mode networks change in the hypothesized opposite directions pre- to post-task performance. Additionally, we identified two subnetworks within the hippocampal network. The primary subnetwork was independent of the sleep condition and increased in functional connectivity pre- to post-task performance and after the break. Connectivity in the second subnetwork decreased over the break only for participants who took a nap. Finally, spindle-slow oscillation coupling power was positively associated with memory performance and decreased connectivity in the second, sleep-dependent hippocampal subnetwork. All effects were the same for allocentric and egocentric training. Our findings suggest that resting-state activations can be divided into task-related and consolidation-related changes that are either sleep-independent or sleep-dependent. Only sleep-dependent changes are associated with fast spindle-slow oscillation coupling that contributes equally to memory performance for allocentric and egocentric learning.

## Methods & Materials

### Participants

Seventy-seven neurologically healthy, right-handed male participants (age range: 18-30 years, mean = 24) were recruited for the study. Only male participants were selected because the sample comprises a subpopulation of a larger translational body of research using male rodents only (Samanta et al., 2021). The study was approved by the local ethics committee (CMO Arnhem-Nijmegen, Radboud University Medical Center) under the general ethics approval (‘Imaging Human Cognition,’ CMO 2014/288), and the experiment was conducted in compliance with these guidelines. Recruitment took place via the Radboud Research Participation System. Participants were screened for and excluded based on 1) taking sleep medications 2) regular naps, and 3) being involved in professional gaming activities. Prior to the start of the experiment, all participants provided written informed consent and received financial compensation for their participation. In total, 19 participants were excluded from the analysis after data acquisition. Eight subjects were excluded from the experiment due to technical issues during learning or retrieval: For five participants, the joystick was incorrectly calibrated, and for the remaining three, there were technical problems including abrupt crashing of the task environment program during the scan. Three subjects slept significantly less than the estimated nap duration of 90 min. For another four subjects large image displacement was identified due to head movement during the fMRI scans. The amount of displacement in the image series was assessed with a quality index (QI) using Eq. (1):

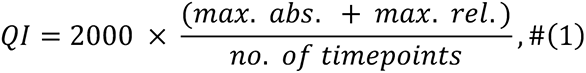

where *max. abs.* is the maximum absolute displacement of a given volume with respect to a reference time point (i.e. middle volume in the time-series), and *max. rel.* represents the maximum relative displacement at a given volume with respect to the subsequent time point. The two measures account for gradual as well as abrupt changes in head position. The cut-off for exclusion of participants was set to a QI > 16 (Ribeiro de Paula et al., 2017).

### Experimental Procedure

With a 2×2 full-factorial mixed design we tested system-wide consolidation processes of a spatial memory paradigm. During fMRI scans, participants engaged in allo- or egocentric spatial learning and memory retrieval in a previously validated human analogue of the water maze (Muller et al., 2018; Schoenfeld et al., 2017). Immediately before and after the learning and retrieval sessions, functional connectivity was assessed with resting-state fMRI. This amounts to a total of four resting-state fMRI sessions of 9 min each. In between spatial learning and memory retrieval, sleep EEG was used to measure sleep. In the experimental condition, participants were instructed to sleep for 90 min, while the control group watched a movie for the same amount of time. Approximately two weeks after the experiment, participants in the sleep condition completed a sleep control session for 90 min, which did not follow a learning experience.

### The Virtual Water Maze Task

The virtual water maze task is an adaptation of the rat water maze, an established method to test spatial abilities in rodents (Morris, 1981). This analogous paradigm for humans is compatible with the MRI-scanner environment (Schoenfeld et al., 2017). Participants were instructed to navigate around a virtual island with the objective of finding a treasure box. We used two different settings in this paradigm: a ‘hidden island’ and a ‘cued island’. The hidden island was surrounded by four landmarks (i.e. a bridge, a sailboat, a wind turbine, and a lighthouse; see Fig. S1). A hidden treasure box on the island marked the target location. The box was located in a small indentation on the virtual island surface and, thus, only visible in close proximity. In contrast, the cued island did not have any landmarks for orientation except for a flag next to a treasure box that was visible from a distance. The position of this flag along with the treasure box changed each trial.

The task sequence consisted of 8 alternating blocks of cued and hidden islands resulting in a total of 16 trials. In each trial, participants were free to move around the island in their own pace using a joystick. When the target location in close proximity of the treasure box was reached, the participants could end the trial with a button press on the joystick. There was a 15-s interval between the end of one trial and the start of the next, during which the participants could turn around on the island and orient themselves. Participants first encountered the cued island in which they had to find the visible flag. This island was used to control for perceptual and motor processing to isolate memory effects in later analysis. For the encounter with the hidden island, the participants were randomly allocated to either of the learning conditions—allocentric or egocentric. The allocentric group started at a different location on the island in every trial and would have to reorient themselves each time to find the target location, thereby promoting use of place navigation. In the egocentric group, they would have the same start location in every trial and hence could rely on a repeated fixed movement to get to the target location in addition the visible cues, allowing them to use both navigational strategies. In general, participants were blinded to these two conditions. For the learning trials, the main objective of the participants in both conditions was to learn the location of the target box across the 8 hidden island trials. For the retrieval trials, the island setup remained the same with one modification—the treasure box was removed from the hidden island and the participants were instructed to mark the location to the best of their knowledge, where they recalled the box to be located.

### Data Acquisition

#### Polysomnographic recordings

Polysomnographic recordings were obtained with a sampling rate of 500 Hz (BrainAmp, Brain Products, Gilching, Germany) during the 90 min naps. 32 scalp electrodes (international 10-20 EEG system) were mounted including Fz, F3, F4, Cz, C3, C4, Pz, P3, P4, Oz, O1, O2 and referenced to the left mastoid. Additionally, horizontal and vertical eye movements (EOG), electromyogram (EMG) on the chin and electrocardiogram (ECG) were recorded. The recordings started shortly after the light was turned off. After the experiment, sleep scoring was performed by an experimenter blinded to the conditions using the SpiSOP toolbox (www.spisop.org; RRID: SCR_015673). In accordance with the American Academy of Sleep Medicine scoring rules (AASM) sleep stages 1 and 2, slow-wave sleep, REM sleep, awakenings, and body movements were visually scored in 30s epochs based on EOG, EMG and the following channels – F3, F4, C3, C4, O1, O2.

#### FMRI acquisition

FMRI time series were acquired using a 3T head-only scanner (Prisma 3T, Siemens, Erlangen, Germany). Seven hundred T2*-weighted images were acquired with a multiband gradient-echo echo-planar imaging sequence (multi-band factor = 8, 64 axial slices, volume repetition time (TR) = 1000 ms, echo time (TE) = 39 ms, 52° flip angle, slice thickness = 2.4 mm; field of view (FOV) 210 mm; voxel size 2mm isotropic). Anatomical images were acquired using a T1-weighted MP- RAGE sequence (192 sagittal slices, volume TR = 2300 ms, TE = 3.03 ms, 8° flip angle, slice thickness, 1 mm; FOV, 256 mm; voxel size 1×1×1 mm).

### Data Processing

#### Sleep EEG preprocessing

The data were preprocessed in Matlab R2020b (Mathworks Inc., Sherbom, MA) using the Sleeptrip toolbox (www.sleeptrip.org; RRID:SCR_017318). All EEG channels were re-referenced to the average mastoids (A1 and A2) and the data were band-pass filtered between 0.3 Hz and 35 Hz (4th-order Butterworth zero-phase filter). In general, only zero-phase filters were applied (i.e. forward and reverse direction). Bad EEG channels were visually rejected based on their power spectra. For the remaining channels, artifacts were detected in 30-s long windows. Segments that were visually identified as body movement were marked for later exclusion. To further identify segments that strongly deviated from the observed overall amplitude distribution, mean amplitude differences for each segment were z-standardized within each channel. Segments with a z > 5 in any of the channels were excluded (Muehlroth et al., 2019). TST was calculated as time spent in stage 1, 2, SWS, and REM sleep. WASO was defined as the time participants were awake between sleep onset and final awakening.

#### Spindle detection

Spindles were detected using an automated algorithm implemented in Sleeptrip based on previous publications (Weber et al., 2020) and the ‘fooof’ algorithm implemented in Python, version 3.8 (Donoghue et al., 2020). Putative power peaks in slow spindle (9-12.5 Hz) and fast spindle (12.5 – 16Hz) frequency bands were visually detected from NREM epochs (i.e., N2 and N3). Before the inspection of the power spectra, the aperiodic 1/f component was removed from the signal by fitting a Lorentzian function. For each participant, slow spindle peaks were manually identified in the average of the 3 frontal channels (Fz, F3, F4; Sleep Test Session: 10.25 Hz ± 0.1, mean ± SEM; Sleep Control Session: 10.07 Hz ± 0.14, mean ± SEM) and fast spindles peaks in the average of the 3 central channels (Cz, C3, C4; Sleep Test Session: 13.44 Hz ± 0.09, mean ± SEM; Sleep Control Session: 13.44 Hz ± 0.06, mean ± SEM). To verify the visually defined frequency peaks, we iteratively fitted Gaussians to the periodic power components with fooof. In case of a difference in the manual labels and the fitted Gaussian peaks, the power spectra were revisited. This method was used to avoid the introduction of spurious spindle events (Ujma et al., 2015). Then, the signal of every channel was band-pass filtered (± 1.5 Hz, − 3 dB cutoff, 4th-order Butterworth zero-phase filter) around the identified individual frequency peaks. The root-mean-square (RMS) representation of the signal was calculated at every sample point and smoothed using a moving average filter in a 200 ms sliding window. A potential spindle was marked if the amplitude of the smoothed RMS signal exceeded its mean by 1.5 SD of the filtered signal for 0.5 to 3 seconds. Local minima and maxima in the filtered spindle signal were marked as peaks and troughs. The largest trough was defined as the spindle peak time. The spindle amplitude was defined by the potential difference between the largest trough and the largest peak. Individual frequency of a concrete spindle was determined by adding the number of all troughs and peaks and dividing by twice the duration of the respective spindle. Spindles with boundaries closer than 0.25 s were eventually merged.

#### Slow oscillation detection

Detection of slow oscillations at frontal electrodes was also based on an automated algorithm implemented in Sleeptrip and validated elsewhere (Ngo et al., 2013). For all NREM epochs, the pre-processed EEG data was low-pass filtered at 4 Hz (6th-order Butterworth zero-phase filter). The whole signal was then divided into negative and positive half-waves that were separated by zero-crossings. A potential slow oscillation was defined as a negative half-wave followed by a positive half-wave at a frequency range between 0.5 Hz and 1 Hz. Slow oscillation amplitude and frequency were computed with the same approach used for spindles. Putative slow oscillations exceeding a trough of 1.25 times the mean trough of all putative slow oscillations as well as an amplitude of 1.25 times the average amplitude of all potential slow oscillations were accepted for further analysis.

#### Resting-state fMRI preprocessing

The MRI DICOM files were entered into an automatic pipeline in GraphICA (BraiNet—Brain Imaging Solution Inc.—Sarnia, ON, Canada). Anatomical and functional images were preprocessed using FSL 6.03. Preprocessing steps of the T1-weighted anatomical images included bias-field correction (RF/B1-inhomogeneity-correction), brain-extraction, tissue-type segmentation (CSF, GM, WM) and subcortical structure segmentation. On the functional data we performed skull stripping, motion correction, slice-timing correction, spatial smoothing, which is calculated based on the voxel size, ceiling (1.5*voxel size), coregistration, ICA-based automatic removal of motion artifacts, high-pass filtering and nuisance regression (WM and CSF). The data were not normalized to a reference space as characteristics of the natural brain space should be preserved in the following analysis.

#### Dual regression analysis

Dual-regression was used to identify subject-specific spatial maps using 11 resting-state network masks: Auditory, Default Mode Network, Executive Control Left, Executive Control Right, Hippocampal, Language, Salience, Sensorimotor, Visual Lateral, Visual Medial and Visual Occipital. The intensity of the component spatial maps was expressed in units of percent signal change from the mean.

#### Regional parcellation

Each subject’s T1-weighted image was automatically segmented with a pipeline implemented in Freesurfer 7 (v7.1.0, http://surfer.nmr.mgh.harvard.edu/). Further parcellation was performed with GraphICA using a gradient-weighted Markov Random Field model procedure described in (Schaefer et al., 2018). This parcellation model contains three competing terms that tradeoff optimal properties of the final segmentation: a global similarity term to group brain locations with similar image intensities, a local gradient term to detect abrupt functional connectivity changes between neighboring brain locations, and a spatial connectedness term to ensure spatial connectedness within parcels. The procedure yielded 832 parcellated brain regions which were included as network nodes for further analyses.

#### Functional network construction & thresholding

Using GraphICA each nodal fMRI time course was correlated with the IC time course for the three resting-state networks separately. After the coregistration of each of the functional resting state network to the subject in native space, a mean z-value was calculated by averaging the scalar map values of the voxel belonging to each one of the 832 ROIs. Resulting *z-*standardized correlation coefficients describe the loading of each nodal time course on the respective resting-state network. To remove spurious or weak *z-*values, for instance due to noise, the loadings were thresholded with a data-driven mixture modeling approach at single-subject level (Bielczyk et al., 2018). Additionally, the thresholded networks were matched to network templates provided by GraphICA that were derived from 200 healthy controls. Only those nodes found in the templates were considered for further analysis (i.e., hippocampal network = 74 nodes, default mode network = 171 nodes, right executive control network = 164 nodes).

### Statistical Analysis

#### Sleep EEG analysis

Statistical analyses of the sleep EEG data were conducted using Matlab R2020b (Mathworks Inc., Sherbom, MA) with the open-source toolbox Fieldtrip and the Circ-Stats toolbox (Berens, 2009). The analysis is confined to channels Fz for slow oscillations and slow spindles and Cz for fast spindles. Not all sleep variables used in our analysis followed a normal distribution. All within group comparisons of the sleep descriptives and oscillation properties between the sleep session after learning and the sleep control session were tested using non-parametric Wilcoxon Signed-Rank Tests for dependent samples. Between-group comparisons of the sleep variables in the allocentric versus egocentric condition were computed with non-parametric Mann-Whitney *U* Tests for independent samples. For these tests median and quartile values of the variables were reported. A corrected p-value < 0.05 was viewed as statistically significant.

#### Temporal relation between detected slow oscillations & spindles

The coordination of slow oscillations and spindles was evaluated for NREM sleep as spindle–slow oscillation coupling has been demonstrated to be stable across stage 2 and slow-wave sleep (Cox et al., 2017) even though the type of slow oscillation is different (K-complexes vs. Delta waves). The general temporal relation between slow oscillation and spindle events was calculated by determining the proportion of spindles whose center (i.e., maximum trough) occurred in an interval of ±1.2 s around the trough of the identified slow oscillations, and the amount of slow oscillations with spindles whose center occurred within ±1.2 s around the respective trough of the oscillation. The time window of ±1.2 s was chosen to cover one whole slow oscillation cycle (0.5–1 Hz, i.e., 1–2 s). Whenever two slow oscillation time windows overlapped the one being further away from the spindle center was removed. The exact timing of slow oscillation and spindle events was visualized by PETHs (Fig. 2b) of fast and slow spindle centers (test events) occurring within a time interval of ±1.2 s around each slow oscillation trough (target event). Probabilities of seed event occurrence were summed within bins of 100 ms and transformed into percentages. To test the temporal pattern of the PETHs, we implemented a randomization procedure by randomly shuffling the order of the PETH bins 1000 times. The resulting surrogates were averaged for each individual and tested against the original PETHs using dependent sample *t*-tests. Control for multiple comparisons was achieved through cluster-based permutation testing with 5000 permutations (Maris & Oostenveld, 2007).

#### Event-locked phase coupling

For event-locked cross-frequency analyses (Helfrich et al., 2018), artifact-free slow oscillation-event trials over channel Fz (±3 s around the maximum slow oscillation trough) were selected and band-pass filtered in the slow oscillation frequency range (0.3 - 1.25 Hz). After applying a Hilbert transform, we extracted the instantaneous phase angle from the analytical signal. The same trials were also band-pass filtered between 9-12.5 Hz (slow spindles component) and between 12.5-16 Hz (fast spindles component) to compute the amplitude envelop from the analytical signal. We only considered the time range from −2 to 2 s to avoid filter edge artifacts. For every subject, channel, and epoch, we now detected the maximum spindle amplitude and corresponding slow oscillation phase angle. Rayleigh tests were computed to test the phase distribution per subject against uniformity. The mean circular direction and resultant vector length across all NREM events were determined. PPC was used as an unbiased estimator of the consistency in phase relationship between spindles and slow oscillations (Vinck et al., 2012) based on the spindle-trough triggered Fourier spectra in a time window of −3.5 to +3.5s (100ms steps) and a slow oscillation frequency range from 0.3 to 3 Hz.

#### Time–frequency analysis

To further describe the temporal relation between spindles and slow oscillations, we again used the previously computed slow oscillation-event trials (6-s segments). These trials were matched with a randomly chosen artefact-free time segment of 6 s during the same sleep stage as the respective slow oscillation used as baseline trials. For the time-frequency analysis of trials with and without slow oscillations we used superlets (Moca et al., 2021). Superlets are composed of a series of Morlet wavelets with the same center frequency but increasingly constrained bandwidth that have higher time-frequency resolution than established approaches. Specifically, we applied a fractional adaptive superlet transform (FASLT) to the preprocessed data of slow oscillation and baseline trials between 5 Hz and 20 Hz in steps of 0.25 Hz, with and initial wavelet order of 3 and an order interval between 5 and 15 (Barzan et al., 2021). The time-frequency representations of slow oscillation-event trials and control trials were then compared for each subject using independent-samples *t*-tests. The resulting *t*-maps reflect the increase/decrease in both the fast and slow spindle frequency range for trials with slow oscillations, compared to trials without. To calculate the within-subject contrasts, we drew 100 random sets of baseline and slow oscillation trials while maintaining the ratio of stage 2 to slow-wave sleep trials. *T*-maps for all these random trial sets were averaged for each subject. Separately for each memory condition, *t*-maps were tested against zero using a cluster-based permutation test with 5000 permutations in a time window of –1.2 to 1.2 s.

#### Resting-state fMRI analysis

fMRI data of the three resting-state networks were analyzed in RStudio 1.4.1103 (RStudio, Inc., Boston, MA). All analyses were performed for the three pre-selected resting-state networks separately (i.e., hippocampal network, default mode network, right executive control network) and were based on the nodal *z-*values obtained per IC that describe the connectivity pattern in the functional resting-state networks. We explored three different approaches to analyze the data including Graph Theory analysis, Euclidean Distance matrices and SVD out of which SVD was most suitable for the current data set. SVD was performed on the raw z-values to capture the original organization of the resting-state networks more accurately. The thresholded *z-*values formed an *m x n* data matrix (*m* = no. of participants x 4 sessions; *n* = no. of nodes). An SVD for sparse matrices was computed as dimensionality reduction since the thresholded data matrix contained mainly zero values. Two principal modes per network were extracted which together explained 15-20% of the variance in the data. Each mode can be understood as a variance-based functional resting-state subnetwork (see Supplementary Movies). As a result of the SVD, we obtained an averaged low-rank reconstruction of the networks as a rank-r approximation that describes the spatial distribution of the respective subnetwork, using Eq. (2):

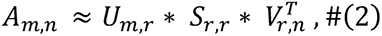

where *A* is the original data matrix that is approximated by *U* containing *r* left singular vectors, S containing *r* singular values and *V* containing *r* right singular vectors. In preparation of statistical testing, the respective left singular vector (u_m,i_) was scaled by its singular value (s_i,i_), where *i* is the *i*-th row/column of the matrix. The result (u_m,i *_ s_i,i_) is a column vector that represents the scores of the participants for all sessions on the *i*-th subnetwork. These values can be interpreted as the connectivity strength of the selected functional subnetwork for all participants, across sessions and conditions. For each of the two subnetwork, the obtained participant scores were analyzed as dependent variables using a linear mixed effects model. Model predictors included sleep condition (sleep, wake), session (1- 4 or pre/post-break, pre/post-task) and memory condition (allocentric, egocentric) as fixed effects. Participants were modelled as random effects for the four resting-state sessions. Akaike Information Criterion (Akaike, 1973), Bayesian Information Criterion (Schwarz, 1978) and log-likelihood were used to assess the model fit.

## Results

### Study Design and Memory Performance

The present study used a one-day paradigm to test consolidation of a spatial memory task (Fig. 1a). During fMRI scans, participants performed allo- or egocentric spatial learning and memory retrieval in a previously validated virtual analog of the water maze for humans (Muller et al., 2018; Schoenfeld et al., 2017). A detailed task description can be found in the first publication of this dataset (Samanta et al., 2021). Briefly, participants (n = 65) freely navigated a virtual island using a joystick to reach a fixed target location. In the learning session, the location was indicated by a hidden treasure box. However, during retrieval, participants were instructed to mark the treasure box location to the best of their knowledge without the box being present. Allocentric and egocentric conditions differed such that the randomly assigned participants began each trial either at the same or a changing starting position (Fig. 1b). Memory performance was measured as latency to reach the target location from the starting point. At retrieval, participants generally performed better in the egocentric condition than in the allocentric condition, and the main effect of sleep on memory performance was marginally significant (Fig. 1c; two-way ANOVA with allo/ego F_1,66_ = 15.66, p < 0.001 and sleep/wake F_1,66_ = 3.34, p = 0.07). Looking back at the results presented by Samanta et al. (2021) who used the same dataset, our sleep effect is less close to reach significance (F_1,69_ = 3.8, p = 0.054 vs. F_1,66_ = 3.34, p = 0.07). This discrepancy is explained by a smaller sample size in our study due to our analysis requirements (cf. Methods and Materials, subjects slept significantly less than 90 min or moved too much during resting state scans), leading to a decrease in statistical power to detect a sleep effect.

**Figure 1.**
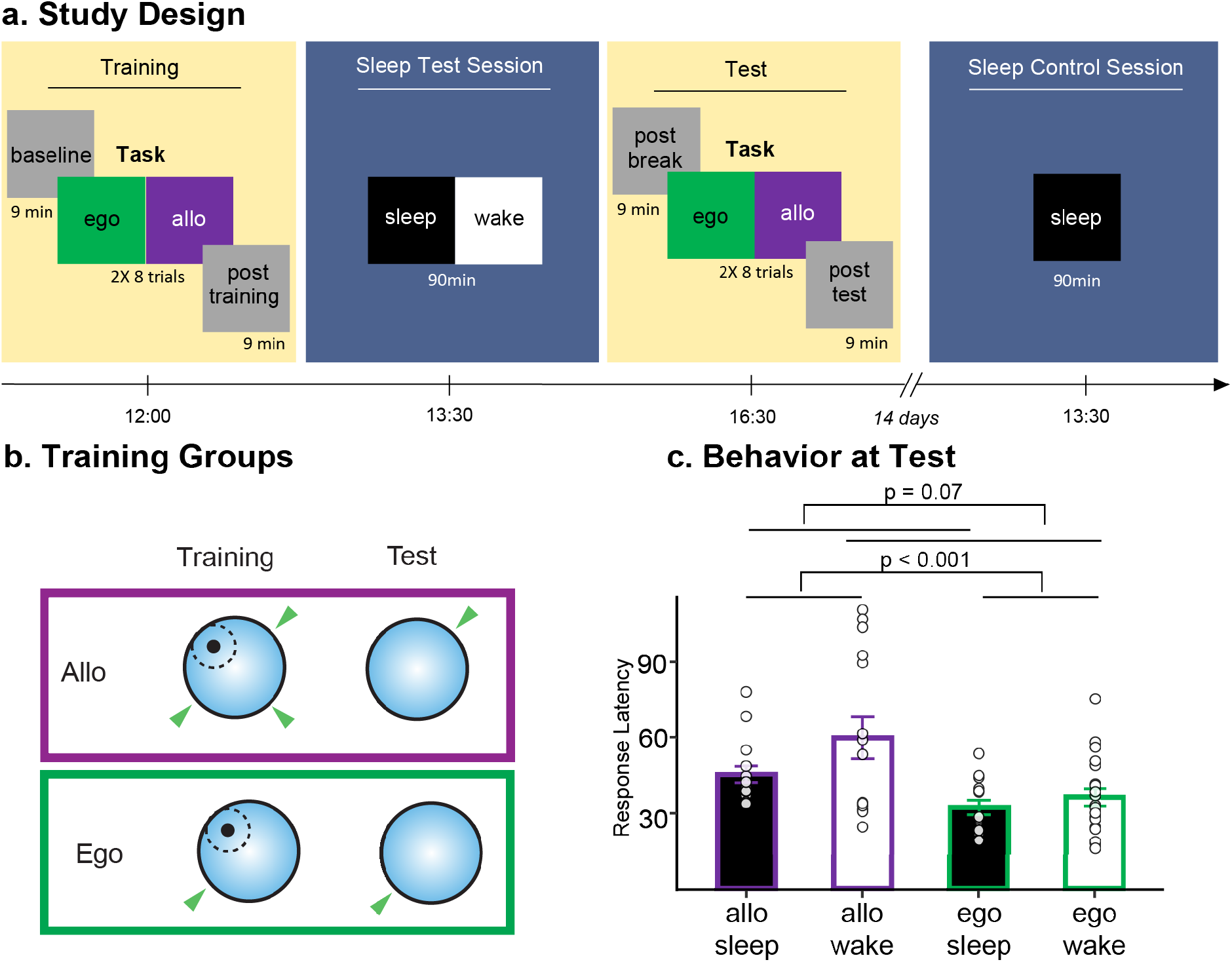
Experimental procedure, task design and memory performance. **a**, A 2×2 full-factorial design with four groups: sleep–allocentric (n = 15), sleep–egocentric (n = 15), wake–allocentric (n = 15), wake–egocentric (n = 20). Subjects participated in two fMRI sessions in which they learned and retrieved a spatial memory task, either in the egocentric or allocentric condition. In both sessions, resting-state fMRI was measured for 9 min before and after task performance. In between the learning and retrieval session, participants were further divided into the sleep or wake control condition, in which they took a nap or watched a movie for 90 min. These measurements were acquired on the same day. Participants who were in the sleep condition returned for a nap control session of 90 min after **∼** 14 days. **b**, Learning-retrieval task configuration. In the egocentric condition participants started from the same island locations on every learning trial. In contrast, in the allocentric condition, the starting location changed for every learning trial. The target location was marked with a treasure box during the learning period but the box was absent in the retrieval session. **c**, Behavioral results obtained at retrieval. Participants in the egocentric condition (purple bar contours) showed a significantly better memory performance compared to participants in the allocentric condition (green bar contours) Lower latency scores indicate better performance. Participants who took a nap (black bar fillings) did not perform significantly better compared to those that stayed awake (white bar fillings). Error bars = SEM.

Immediately before and after the learning and retrieval sessions, functional connectivity was assessed with resting-state fMRI. This amounts to a total of four resting-state fMRI sessions of 9 min each. In between spatial learning and memory retrieval, polysomnography was used to measure sleep architecture (Supplementary Table T1). In the experimental condition, participants slept around 90 min (93.48 ± 1.77, mean ± SEM), while the control group watched a neutral movie for the same amount of time. Approximately two weeks after the experiment, participants in the sleep condition completed a sleep control session for around 90 min (103.30 ± 2.66, mean ± SEM), which did not immediately follow a spatial learning experience. The sequence of the two sleep sessions was not counterbalanced to keep subjects blinded to the experimental conditions at the test session. Using a similar task design, Genzel et al. (2014; 2012) obtained comparable nap durations. However, here, participants slept significantly longer in the sleep control session than in the sleep test session (Wilcoxon Sign-Rank Test, z = −2.68, p = 0.01). This session effect hints at a habituation to the sleep laboratory. The difference to previous studies may be due to the use of a different laboratory setting, for Genzel et al (2014; 2012) the sleep laboratory (Max Planck Institute of Psychiatry) was designated as such, while the sleep laboratory at the Donders Institute is the EEG lab (and therefore less “homely”).

### Sleep Properties at Test and Control Session

Spatiotemporal mechanisms during NREM sleep have been shown to consolidate experiences encoded during prior wakefulness (Maingret et al., 2016). At the systems level, experiences are thought to be reactivated in the hippocampo-thalamo-cortical network during sleep to form long-term representations of these experiences. Various oscillations involved in memory consolidation may coordinate the communication between these brain structures (Genzel, 2020; Genzel, Kroes, et al., 2014). We were interested in slow oscillations, as well as slow spindles (9–12.5 Hz) and fast spindles (12.5–16 Hz) during NREM sleep, which may play a distinct role in the process of memory consolidation (Genzel & Robertson, 2015). In line with the previously reported topography of these sleep oscillations in human EEG (Adamczyk et al., 2015), we focused our analyses on frontal slow oscillations and slow spindles (i.e., channel Fz) and central fast spindles (i.e., channel Cz). To capture specific influences of the spatial learning experience on the sleep architecture of the sleep test session, we compared sleep descriptives as well as spindle and slow oscillation properties (see Supplementary Table T1) between the sleep test and control sessions.

We found that participants slept significantly longer in the control session compared to the test session. In comparison to the test session, also fast spindle amplitude was significantly higher in the control session (Wilcoxon Sign-Rank Test, z = −2.91, p < 0.01). We further evaluated whether the significant difference in total sleep time (TST) between test and control sessions could predict this difference in fast spindle amplitude, controlling for wakefulness after sleep onset (WASO). However, there was no significant effect of TST on fast spindle amplitude differences across sleep sessions (multiple linear regression, TST: β = 0.529, p = 0.601). The increase in fast spindle amplitude, similar to the increase in TST, could be due to an uncontrolled habituation effect on the sleep laboratory. Nevertheless, we hypothesized the temporal relationship of sleep oscillations to be crucial for memory consolidation. In this respect, we did not observe any differences between sessions, as will be presented in the following.

### Spindle-Slow Oscillation Coupling at Test and Control Session

The coupling between sharp wave-ripples, spindles and slow oscillations may set a crucial time window for consolidating a learning experience (Genzel, Kroes, et al., 2014). Therefore, we focused on the temporal relationship between spindles and slow oscillations to further explore a difference in sleep micro-architecture between sleep test and control sessions. First, we computed the number of spindles’ maximum troughs (in %) occurring in one cycle (interval of ± 1.2 s) around the center of the identified slow oscillations (Fig. 2a). In both sessions we detected similar number of fast spindles coupled to slow oscillations (Wilcoxon Sign-Rank Test, z = −1.14, p = 0.26) and slow spindles coupled to slow oscillations (Wilcoxon Sign-Rank Test, z = −0.75, p = 0.45). Reversely, roughly equal number of slow oscillations were coupled to fast spindles (Wilcoxon Sign-Rank Test, z = 1.02, p = 0.31) and slow spindles (Wilcoxon Sign-Rank Test, z = 1.10, p = 0.27) following learning and at sleep control. These findings suggest that spindles and slow oscillations are equally bound to each other. Generally, we detected a consistent spindle-slow oscillation coupling across both sleep sessions independent of the spatial learning experience conducted immediately before the sleep test session.

**Figure 2.**
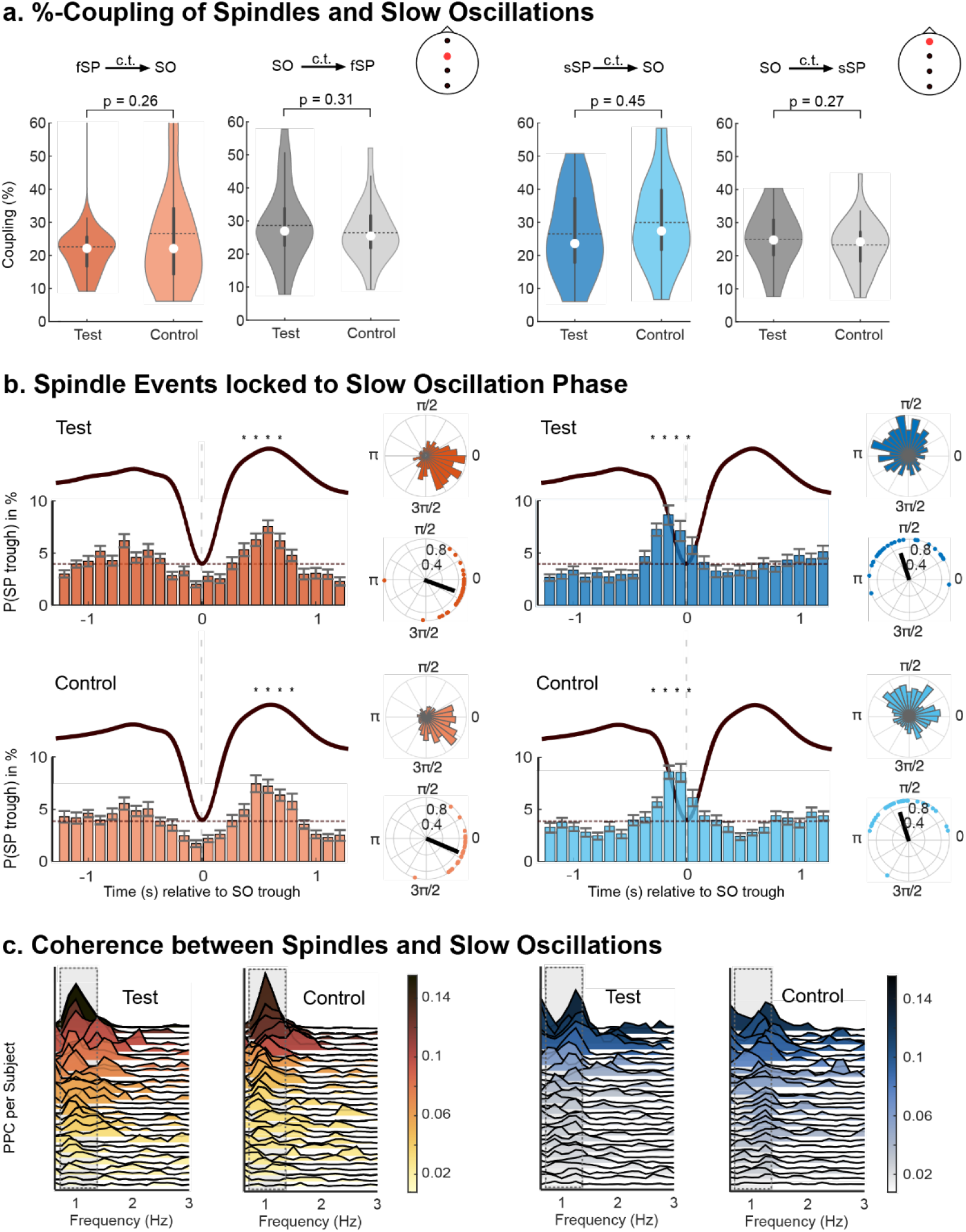
Spindle-slow oscillation coupling across sleep test and control sessions. SO: slow oscillation; sSP: slow spindle, fSP: fast spindle, Test: sleep test session, Control: sleep control session. **a**, %-Coupling of spindles to slow oscillations and vice versa. Fast and slow spindles coincide with slow oscillations in both sleep sessions. About one fourth of all fast spindles (orange) and slow spindles (blue) occur within an interval of ±1.2 s around the slow oscillation trough. Approximately an equal amount of slow oscillations is coupled to the occurrence of spindles (grey) in the same interval. Violin plots show the median (white dot), mean (dotted line), IQR and sample distributions. c.t.: ‘coupled to’. **b**, Specific Time and Phase relationship between spindles and slow oscillations. PETHs of slow and fast spindles co-occurring with frontal slow oscillations are depicted for both sleep sessions. The reference distribution obtained after randomization of the data is shown by horizontal dashed line. Asterisks indicate significantly increased spindle occurrence contrasted with the reference distribution (cluster-based permutation test, cluster α < 0.05, positive clusters only). Vertical dashed lines mark the slow oscillation trough. Average Slow-Oscillation ERPs are shown for each session. In both sessions, frontal slow spindles peak at the up- to down-state transition before the trough (significant positive cluster: −300–100 ms). Central fast spindles prominently peak during the slow oscillation peak (Test: significant positive cluster: 400–700 ms, Control: significant positive cluster: 500–800 ms). Error bars of 100-ms time bins = SEM. Polar plots show spindle-slow oscillation coupling for one example subject (top) and group level results (bottom). Top: Polar histograms display maximum spindle amplitude per slow oscillation phase. Note the peak in the right lower quadrant for fast spindles (i.e. 3π/2 – 2π) and the left upper quadrant for slow spindles (i.e. π/2 – π). Mean slow oscillation phase with sleep spindle power peaks. Bottom: Dots depict individual subjects and the black line the average of the sample results. **c**, Consistency in phase relationship between spindles and slow oscillations. Ridge-line plots of fast spindle-PPCs (orange) and slow spindles-PPCs (blue) for all subjects across sleep sessions. PPC values are expressed by color and height of the ridgelines. Slow oscillation frequencies (0.3 – 1.25 Hz) are outlined with black dotted line.

Successful memory consolidation may depend on a coupling precision of tens or even hundreds of ms (Maingret et al., 2016; Muehlroth et al., 2019). Thus, we analyzed the precise spindle-slow oscillation coordination using peri-event time histograms (PETHs; Fig. 2b). The PETHs show the likelihood of maximum spindle-troughs occurring in the ± 1.2 s time-window around the slow oscillation events within 100-ms time bins. Fast spindles co-occurred immediately before and at the slow oscillation peak following the slow-oscillation trough (cluster-based permutations: sign. cluster: 400-700 ms at test, p < 0.001; 500-800 ms at control, p < 0.001). On the other hand, slow spindles coincided in transition to and with the slow oscillation trough (cluster-based permutations: both sign. clusters at −300-100 ms, p < 0.001). The robustness of this effect was tested against a reference distribution obtained by randomly shuffling the time stamps of the spindle troughs. The temporal specificity of spindle-slow oscillation coupling did not differ across the two sleep sessions, again suggesting the presence of a general coupling mechanism independent of a spatial learning experience.

Previous findings have demonstrated higher spindle-slow oscillation coupling strength for young compared to old adults, and following gross motor learning (Cross et al., 2020; Helfrich et al., 2018). We were interested in elucidating this consistency in phase relationship in the context of spatial learning. Polar plots in Fig. 2b show the coupling phase for one example subject and the whole sample, computed as the instantaneous slow oscillation phase (0.3-1.25 Hz) at the time of the maximum spindle amplitude over channel Fz for slow spindles (9-12.5 Hz) and channel Cz for fast spindles (12.5-16 Hz). Already at the single-subject level, a precise spindle-slow oscillation coupling can be seen, which is different for fast spindles and slow spindles but relatively consistent across sleep sessions (see polar histograms, Fig. 2b). At the group level, most fast spindles occurred between 5.76-6.28 radians of the slow oscillation (Test: 6.03 ± 0.56, Control: 6.07 ± 0.33, circular mean ± std), whereas slow spindles occurred between 1.57-2.62 radians of the slow oscillation (Test: 1.89 ± 0.79, Control: 1.87 ± 0.71, circular mean ± std; Fig. 2e). These findings match the temporal coordination between spindles and slow oscillations shown in the PETHs.

To estimate the consistency of these phase relationships we calculated pairwise phase consistency (PPC; cf. Methods and Materials). The maximum PPC values neither differed for fast spindles nor for slow spindles between the two sleep sessions (Wilcoxon Sign-Rank Tests, fast spindle max PPC: z = 0.451, p = 0.652, slow spindle: max PPC: z = −0.067, p = 0.947). Additionally, we observed noticeable individual differences in PPC (Fig. 2c). We were interested whether this variability in PPC was related to the participant’s general sleep quality (measured with PSQI), which was not significant (linear correlation, fast spindles: ρ = 0.30, p = 0.14, slow spindles: ρ = 0.15, p = 0.46).

In summary, spindles were consistently coupled to slow oscillations with a specific phase relationship that supports previous findings in the time domain. The overall coupling results presented so far did not differ across the two sleep sessions. As described earlier, we observed a longer total sleep time in the sleep control than the sleep test session, which might have had compensatory effects on the sleep architecture behind which a difference in coupling was hidden. Thus, we present the following results for the sleep test session only.

### Fast Spindle-Slow Oscillation power is Associated with Memory Performance

Having established general time and phase dynamics of spindle-slow oscillation coupling across sleep sessions, we proceeded our analysis with a focus on the two memory conditions (allo vs. ego) during the sleep test session as sleep may have a distinct effect on allocentric versus egocentric spatial memories. Possible differences between the groups were tested for %-coupling and PPC (Fig. 2a and Fig. 2f). Neither the amount of spindle-slow oscillation coupling nor the consistency in spindle-slow oscillation phase was significantly different between groups (Mann-Whitney U Test, % fast spindle: z = −1.12, p = 0.27, % slow spindles: z = 0.09, p = 0.93; max. PPC fast spindles: z = 1.742, p = 0.081, max. PPC slow spindles: z = 0.083, p = 0.934). We further probed this pattern by comparing the oscillatory power in the spindle frequency range for time segments with and without slow oscillations (cf. Materials and Methods). Slow oscillation trials were constructed by epoching ± 1.2s around randomly selected slow oscillations. In these trials, the power in the spindle frequency range indeed differed significantly from randomly selected intervals without slow oscillations (for all t-value clusters: p < 0.001, Fig. 3a). This effect was equally present in both memory conditions as the difference map did not contain any significant modulation clusters (Fig. 3a). To conclude, none of the metrics of spindle-slow oscillation coupling differed between memory conditions. The results were also in line with the non-significant interaction between training and sleep for the behavior. Thus, we decided to collapse over the two memory conditions to increase statistical power for further testing.

**Figure 3.**
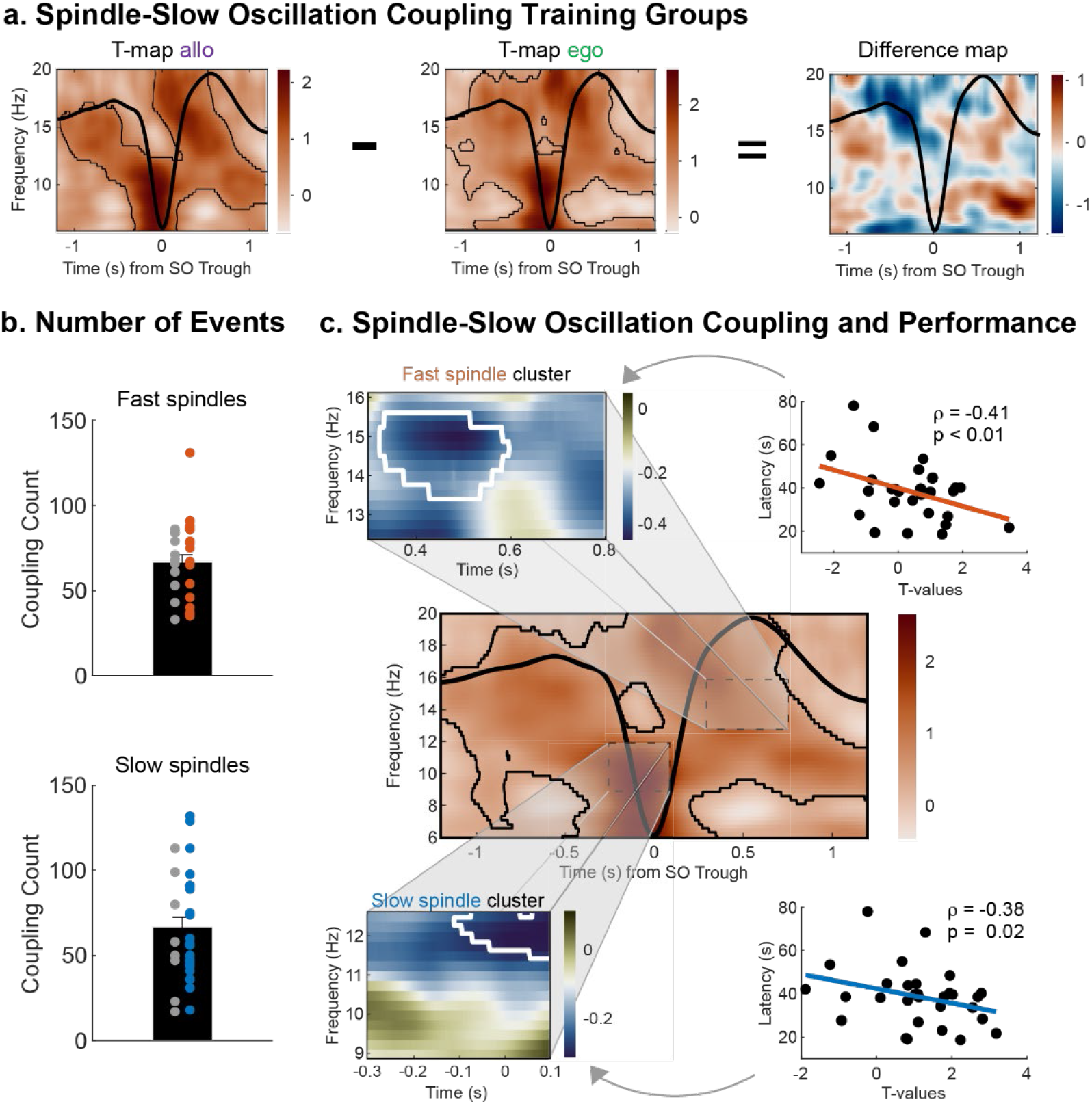
Spindle-slow oscillation coupling at sleep test session and its association with memory performance. **a**, Power modulations in the two memory conditions at sleep test session. Differences in power for slow oscillation trials (trough ± 1.2 s) compared to baseline trials without slow oscillations are depicted (in t-score units) for the allo and ego group separately. T-map ego was subtracted from t-map allo to obtain the difference map. The average frontal slow oscillation for each memory group is overlaid in black. In both groups, EEG activity is modulated as a function of the slow oscillation phase but there is no difference between the two groups. Significant clusters are outlined in black (cluster-based permutation test, cluster α < 0.05) **b**, Bar graphs including sample distribution of coupling count for fast spindles (*top*) and slow spindles (*bottom*) at sleep test session. Colored dots mark the subjects whose spindle-slow oscillation coupling was significantly above chance level (permutation test, α < 0.05). Error bars = SEM. **c**, Correlation of spindle-slow oscillation power modulations and memory performance. *middle:* T-map as outlined in (a) but including all subjects (slow spindles and fast spindle reference windows for later analyses highlighted by dashed black line). *top left*: fast spindle window with significant correlation clusters between slow oscillation-specific EEG activity and memory performance obtained by contrasting the correlation for each time–frequency point against a reference distribution of bootstrapped EEG–behavior correlations. Significant correlation cluster is outlined in white. *top right:* Maximum correlation pixel extracted from the fast spindle cluster contrasting t-statistics against response latency in the memory task. *bottom left:* slow spindle window with correlation values between slow oscillation-specific EEG activity and memory performance obtained as for fast spindle window. *bottom right*: Same as top right but for slow spindles related t-statistics.

Considering the whole sample at the test session, we first checked whether the amount of spindle-slow oscillation coupling was merely a random byproduct of the occurrence of spindle and slow oscillation events. The obtained amount of coupling differed significantly from a random sampling distribution in 60% of the sample (Fig. 3b, permutation tests: p < 0.05, 1000 iterations per participant). Next, we again computed power modulations during slow oscillation trials (Fig. 3c). We found that power modulations during the slow oscillation trials differed significantly from matched control trials in the spindle frequency ranges and the previously established time intervals (see PETHs, Fig. 2b). Next, we performed correlational analyses between slow oscillation–control power differences (expressed in t-values) and memory performance separately for each time–frequency point in the respective spindle cluster extracted from the t-map (Fig.3c). We found significant associations by testing against a bootstrapped reference distribution of the EEG–behavior correlations. In this way, we could identify significant correlation clusters representing the relation between memory performance and a specific pattern of EEG activity modulated during the slow oscillation up-state (cf. Materials and Methods). A significant negative correlation cluster was identified for fast spindles (cluster p < 0.01, max ρ = −0.41, p < 0.01, Fig. 3c top). Similarly, we identified another significant negative correlation cluster for slow spindles (cluster p = 0.018, max ρ = −0.38, p = 0.02, Fig. 3c bottom): more fast-spindle power and slow-spindle power during the slow oscillation period was associated with better memory performance (i.e., lower response latency). These effects were specific for the sleep test session as it could not be observed for the fast spindle or slow spindle cluster in the sleep control session (Supplementary Fig. S3).

To conclude, we found evidence for an association between the coordination of slow oscillations and spindles with memory performance that was specific for sleep after learning in contrast to a control sleep session.

### Functional Resting-State Networks Change across Sessions

After sleep-EEG analysis, we next focused on resting-state functional connectivity before and after learning, consolidation, and retrieval of the memory task to investigate memory consolidation at the network level across resting states. For this, we included fMRI data of the four resting-state sessions (i.e., 1 = baseline, 2 = post-learning, 3 = post-break, 4 = post-retrieval) into the analysis. We were specifically interested in the right executive control network, the default mode network, and the hippocampal network. Task-fMRI analyses of this dataset by Samanta et al. (2021) revealed that regions of these networks changed activation across the consolidation period when participants took a nap but not when they stayed awake. Thus, we obtained the three resting-state networks with GraphICA (Ribeiro de Paula et al., 2017), a tool for resting-state network analysis. For the purpose of dimensionality reduction, we computed the primary modes of variance for each network using singular value decomposition (SVD; cf. Methods and Materials). Finally, we tested the effects of session, sleep conditions, and memory conditions on each participants’ score on the previously identified mode. With this approach we could determine (1) how many functional subnetworks were contained within each network and (2) how the functional connectivity of the networks changed for any of our experimental variables.

For the right executive control network, we extracted a primary subnetwork that was centered on the right inferior parietal sulcus, right inferior parietal lobe, and right dorsolateral prefrontal cortex. In this subnetwork, we found an increase in functional connectivity across task performance in the learning and retrieval sessions (Fig. 4a; linear mixed-effects model, sleep/wake: t_62_ = −0.747, p = 0.458 allo/ego: t_62_ = −0.477, p = 0.635, pre-break/post-break: t_193_ = 1.676, p = 0.095, pre-task/post-task: t_193_ = 2.808, p = 0.005). This increase was driven by change from post-break to post-retrieval (t_193_ = 2.468, p = 0.016). In contrast, the primary default mode subnetwork with main involvement of the posterior cingulate cortex generally decreased in connectivity from pre- to post-task performance and its connectivity was generally higher for the sleep compared to the wake condition (Fig. 4b; linear mixed-effects model, sleep/wake: t_62_ = −2.616, p = 0.011 allo/ego: t_62_ = 0.427, p = 0.671, pre-break/post-break: t_193_ = 1.726, p = 0.084, pre-task/post-task: t_193_ = −2.126, p = 0.034). The decrease in functional connectivity can be explained by a significant change from post-break to post-retrieval (t_193_ = −2.451, p = 0.017). However, the sleep main effect cannot be explained by any of the experimental manipulations and may have arisen by chance (Supplementary Fig. S4). Notably, the primary hippocampal subnetwork which mainly involved the hippocampus and the bilateral parahippocampal cortex showed similar connectivity changes as the right executive control network. In this case, we not only detected a pre- to post-task increase but also a pre- to post-break increase in functional connectivity (Fig. 4c; linear mixed-effects model, sleep/wake: t_62_ =- 0.143, p = 0.887 allo/ego: t_62_ = 0.702 p = 0.485, pre-break/post-break: t_193_ = 3.749, p < 0.001, pre-task/post-task: t_193_ = 4.535, p < 0.001). This was reflected by a significant difference between baseline and post learning (t_193_ = 3.586, p < 0.001) as well as post-break and post-retrieval (t_193_ = 3.338, p < 0.001). Lastly, we tested whether the post-break to post-retrieval change in functional connectivity observed in all networks was associated with memory performance at retrieval. This effect was not significant for the right executive control subnetwork (linear regression, β = 0.016, p = 0.901), the default mode subnetwork (linear regression, β = 0.655, p = 0.422) or the hippocampal subnetwork (linear regression, β = 0.596 p = 0.443).

**Figure 4.**
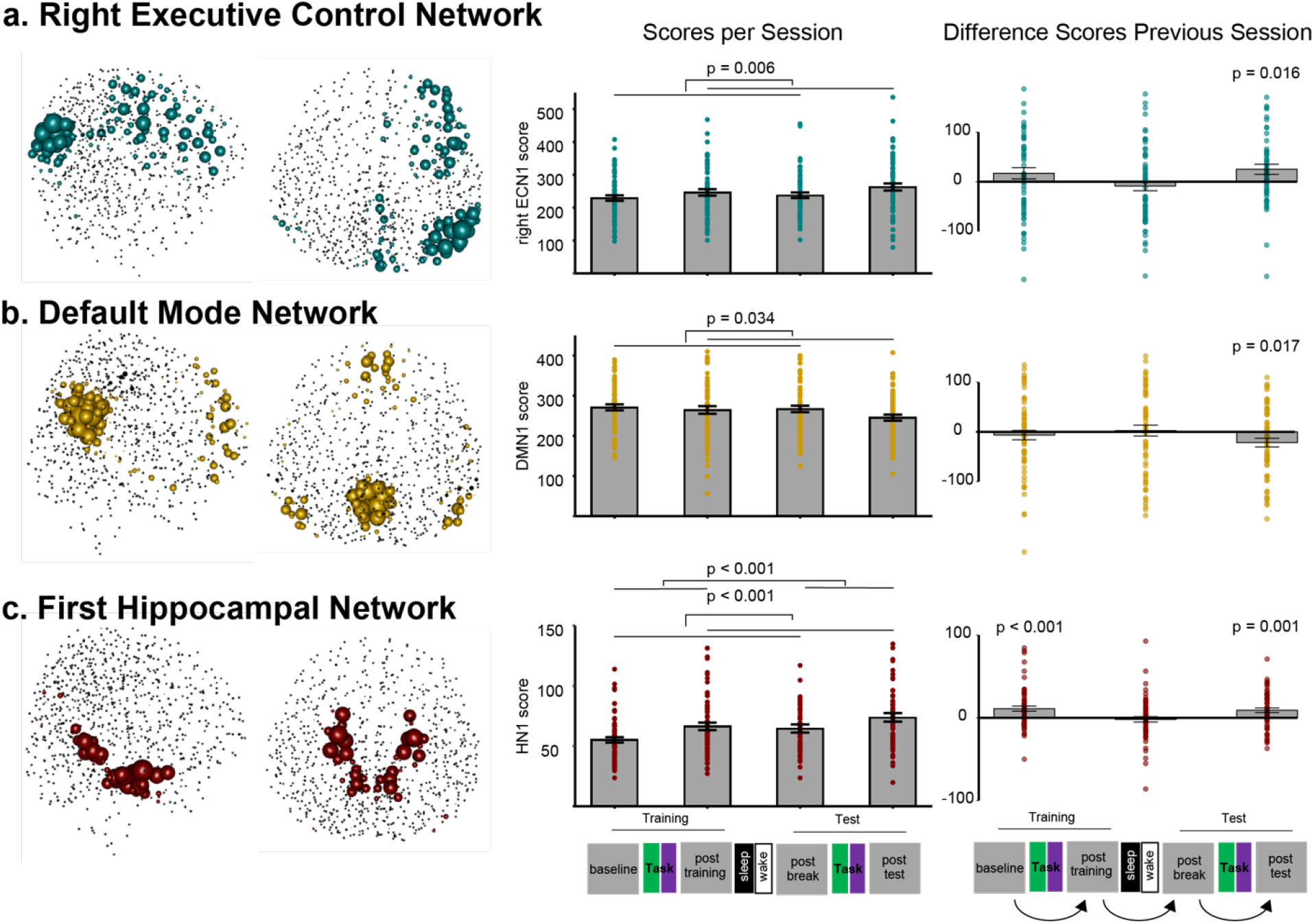
Changes in primary modes of the resting-state networks across sessions. ECN1 = primary executive control network, DMN1 = primary default mode network, HN1 = primary hippocampal network. **a**, *left:* Sagittal and axial representation of the primary mode in the right executive control network. The degree of each one of the 832 regions is represented by the node’s size. *middle:* Results of a linear mixed-effects model with session as within-subject factor, sleep and memory as between-subjects factor and subject*session as random factor. Functional connectivity increased pre- to post-task performance. *right*: Results shown as difference from previous session to next session. Only post-break to post-test showed changes greater than 0. **b**, *left:* Same as in A but for the default mode network. *middle:* Same model as in A. In contrast, the results show that the functional connectivity decreased pre- to post-task performance. *right*: Results shown as difference from previous session to next session. Only post-break to post-test showed changes greater than 0 **c**, *left:* Same as in **a & b** but for the hippocampal network. *middle:* Same analysis as in **a & b**. Note that the functional connectivity increased pre- to post-task performance and pre- to post-break. *right*: Results shown as difference from previous session to next session. Both baseline to post-training and post-break to post-test showed changes greater than 0. Error bars = SEM.

Overall, we consistently obtained a pre- to post-task effect in all three subnetworks, with an increase in the executive control and hippocampal network and a decrease in the default mode network. Only the hippocampal subnetwork increased connectivity pre- to post-break and a sleep main effect was observed for the default mode network. As was shown for our sleep analysis, the factor memory (allo vs. ego) did not impact the changes in the resting-state networks.

### A Sleep-dependent Hippocampal Subnetwork & Fast Spindle-Slow Oscillation Coupling

From the analysis performed by Samanta et al. (2021), it is evident that activation of the precuneus/retrosplenial cortex decreases after sleep but not when participants stayed awake during the consolidation period. Similarly, we extracted a second functional subnetwork of the hippocampal network centered on the bilateral retrosplenial cortex and the bilateral amygdala (Fig. 5a top). We identified this subnetwork based on the SVD factorization, using a singular value spectrum criterion (cf Methods and Materials). Here, we also investigated how session, sleep conditions, and memory conditions affected the participants’ scores on this second mode. In contrast to the task and break main effects detected for the other networks, we now observed an interaction between session and sleep conditions. The subnetwork’s functional connectivity decreased significantly only for participants who took a nap during the consolidation period (linear mixed-effects model, t_191_ = 2.086, p = 0.038). This effect remained when comparing a change from post-learning to post-break sessions only (paired-sample t-test, t_95_ = 2.3971, p = 0.018). Thus, we also computed difference values from one session to the next and tested if these were significantly different from 0. Only significant change occurred for the sleep group from post-training to post-break (t_26_= −3.17, p = 0.004), this was not seen for the wake group (t_38_ = −1.363, p = 0.181).

**Figure 5.**
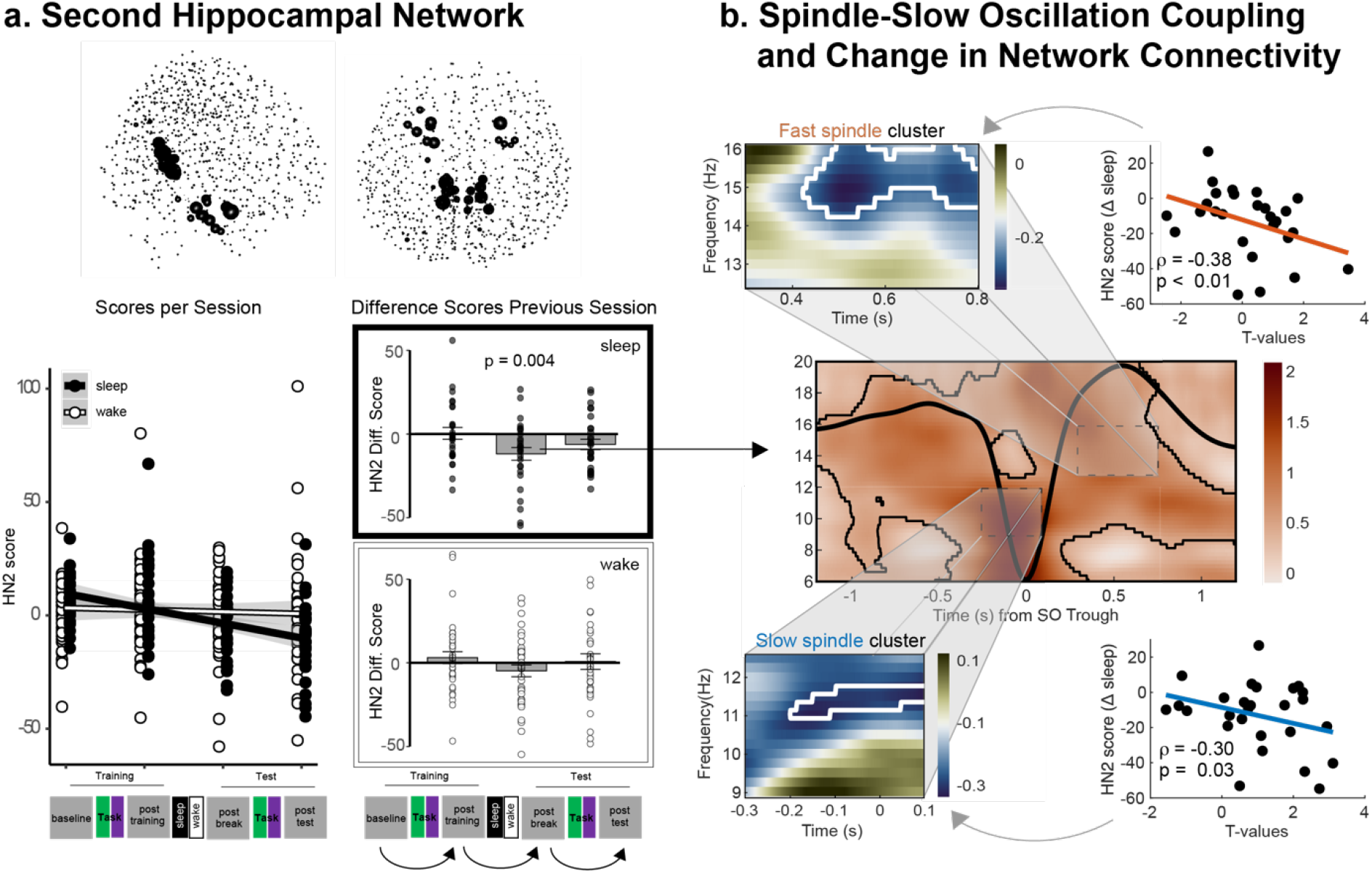
Sleep-related changes in the second hippocampal subnetwork and its association with fast spindle-slow oscillation coupling. **a**, HN2 = second hippocampal network. *top:* Graphical representation of the secondary mode in the hippocampal network. The degree of each one of the 832 regions is represented by the node’s size. *bottom:* Results of a linear mixed-effects model with session as within-subject factor, sleep and memory as between-subjects factor and subject*session as random factor. Functional connectivity decreased significantly in this hippocampal subnetwork across sessions in the sleep condition but remained largely stable in the wake condition. On the right the difference scores are presented. Only for sleep but not wake was there a significant change across the break period. Error bars and shadings = SEM. **b**, *middle:* T-map as outlined in Fig. 3c (slow spindle and fast spindle reference windows for later analyses highlighted by dashed black line). *top left:* Fast spindle window with significant correlation clusters between SO-specific EEG activity and post-learning to post-break change in the hippocampal subnetwork obtained by contrasting the correlation for each time–frequency point against a reference distribution of bootstrapped EEG–behavior correlations. Significant correlation clusters are outlined in white. *top right:* Maximum correlation pixel extracted from the fast spindle cluster contrasting t-statistics against change in secondary hippocampal mode score. *bottom left:* Same as top left but for slow spindle cluster. *bottom right:* Same as top right but for slow spindle related t-statistics.

Therefore, we were interested in whether this decrease in connectivity between post-learning and post-break sessions in the sleep condition was related to the spindle-slow oscillation coupling described previously (see Fig. 3c). Again, correlational analyses were performed – but this time for slow oscillation-control power differences (expressed in t-values) and the change score on the hippocampal subnetwork between the post-learning and post-break session. The correlations were computed for each time– frequency point in the respective spindle clusters extracted from the t-map. A significant negative correlation clusters was identified for fast spindles (cluster: p < 0.001, max pixel: ρ= −0.38, p < 0.01; Fig. 5b, top) and for slow spindles (cluster: p = 0.018, ρ= −0.30, p = 0.03): higher spindle power during the slow oscillation up-state was associated with a larger decrease in functional connectivity of the hippocampal subnetwork across sleep. The clusters partially overlapped with the significant response correlation clusters presented earlier (see Fig. 3c). These findings provide evidence for a relation between the coordination of slow oscillations and spindles and connectivity decrease in the hippocampal subnetwork. They parallel the results obtained for spindle-slow oscillation coupling power and behavior. Therefore, we aimed to establish a triad relationship between change score in this hippocampal subnetwork, spindle-slow oscillation coupling, and memory performance. We regressed the networks’ change score across sleep on memory performance at retrieval. However, the connectivity decrease in the hippocampal subnetwork did not significantly predict behavior (linear regression, β = 0.01, p = 0.94). In summary, we found a hippocampal subnetwork that decreased in functional connectivity across resting-state sessions for participants who took a nap. The functional connectivity remained largely stable for those participants who stayed awake. This effect did not differ between the two memory conditions. The decrease in functional connectivity from post-learning to post-break was positively related to spindle-slow oscillation coupling during sleep. We thus anticipated that this connectivity change would be a correlate of memory consolidation. However, unlike the spindle-slow oscillation coupling power (see Fig. 3c), the sleep-dependent connectivity decrease in the second hippocampal subnetwork did not predict memory performance at retrieval.

## Discussion

This study used resting-state fMRI and sleep EEG data to investigate spatial memory consolidation during a nap or the same period spend awake. Subjects learned a goal location in a virtual watermaze with either different starting locations (allocentric group) or with the same starting location (egocentric group). The sleep-EEG analysis was focused on the spindle-slow oscillation coupling, separating spindles into fast and slow subtypes. Resting-state fMRI was focused on changes in connectivity within previously defined networks. We could show that spindle-slow oscillation coupling correlated positively with memory performance after a nap and also with a decrease in connectivity in a functional hippocampal network including the retrosplenial cortex. This decrease in connectivity was not seen in the group that stayed awake. Task-related increases in connectivity were observed in the executive control network and the other hippocampal network, while a decrease was found in the default mode network. These changes were not specific to the sleep group but seen across all participants.

We replicated previous oscillation coupling findings, showing that slow spindles are nested in the transition period from slow oscillation up- to down-state, and fast spindles frequently occur immediately before and within the slow oscillation up-state. We found no difference in spindle-slow oscillation coupling between allocentric and egocentric training groups. The precise coordination between spindles and slow oscillations was consistently observed across the time, phase, and time-frequency domains. Surprisingly, these effects did not differ between the sleep test session, that immediately followed the spatial learning experience, and the sleep control session, where there was no explicit learning beforehand. The fact that we did not observe a difference between sleep sessions is most likely due to methodological limitations of the study design – the control nap was always the second nap – since also a difference in total sleep time was observed.

The resting-state fMRI analysis was centered on the hippocampal network, default mode network, and right executive control network. These networks were defined and later divided into subnetworks using a data-driven approach. We found brain-wide changes across resting-state sessions that matched our previous findings in the activity analysis (Samanta et al., 2021). Task-related changes in connectivity in the default mode network (decrease in connectivity) opposed changes in the right executive-control network (increase in connectivity). Interestingly, the hippocampal network could be divided into two subnetworks: a primary functional subnetwork centered on the hippocampus and the parahippocampal cortex, and a second functional subnetwork with the retrosplenial cortex as hub region. Functional connectivity in the primary hippocampal subnetwork increased pre- to post-task and pre- to post-break. Functional connectivity in the second hippocampal subnetwork decreased across the four resting-state sessions – most prominently across the sleep break – only for participants who slept during the break but not those that stayed awake. Like our behavior and sleep analysis results, the memory conditions (allo vs. ego) did not affect changes in any of these networks. Our findings suggest task-related and consolidation-related connectivity changes in the functional networks which may be related to memory reactivations.

Finally, we aimed to relate our sleep EEG results to the changes in resting-state connectivity and memory performance. Spindle-slow oscillation coupling was positively related to sleep-dependent changes in the second hippocampal subnetwork and to memory performance. This was specific for sleep after learning and not seen in the control sleep session. Taken together, precise spindle-slow oscillation coupling seems to have behavioral relevance and potentially has functional significance in memory consolidation. This coupling may be a mechanism by which functional connectivity in the hippocampal network is modulated during memory consolidation.

### Connectivity changes in resting state networks

Generally, it has been shown that tasks can affect post-task resting-state brain activity (Lewis et al., 2009). For instance, modulation of learning-dependent spontaneous brain activity after the task has been observed for cognitive tasks involving working memory, emotion, and motor training (Lewis et al., 2009). Such modulation of brain regions associated with a preceding task can be understood as a form of resting-state consolidation, possibly involving similar mechanisms as sleep-related memory consolidation (Tambini & Davachi, 2019). Resting-state connectivity analysis is based on the correlation of different voxels within a network. Neuronal memory reactivations and strengthening of memory networks should lead to increased coactivation of the memory network, which therefore could potentially increase intra-network connectivity as we measure here. Thus, one could interpret our results as changes of reactivations that are occurring during the resting-state.

We show opposite changes of the executive control and main hippocampal network in contrast to the default mode network pre- to post-task performance. The executive control and hippocampal network increased, whereas the default mode network decreased in functional connectivity. If these connectivity changes are induced by occurring reactivations, the results would indicate that after a task one has more reactivations in both the executive control network and the hippocampus. We also showed a sleep-specific decrease in connectivity in a second hippocampal network including the retrosplenial cortex. Potentially this network could be the link between the hippocampus and downstream areas such as parietal cortex, which is part of the executive control network (Genzel, 2020). A decrease in connectivity in this network would therefore potentially indicate that reactivations occurring in the executive control network and the hippocampus would be more independent from each other after sleep. In contrast, after wake when there is no such decrease in connectivity reactivations in the cortex and hippocampus would still be linked.

It has previously been proposed that once consolidation from hippocampus to cortex has progressed, reactivation in the cortex occur independently from hippocampus reactivations. Our results could potentially support this idea. However, the measure of resting-state, within-network connectivity is very indirect and therefore this finding should be confirmed with more direct measure such as simultaneous hippocampal and cortical neuronal recordings in rodents.

### Task-Related Changes in the Right Executive Control and Default Mode Network

Our findings align with the task analysis of this data set performed by Samanta et al. (2021). When measuring changes in BOLD activity across learning and retrieval sessions while participants performed the task, we observed a sleep-related increase in executive control regions and a decrease in default mode regions. In the current analysis, we observed increases of within-network connectivity for the executive control network and decreases in the default mode network over the different resting state session. Of note, the current results, are not specific for the sleep condition in contrast to the activity analysis results from Samanta et al. This discrepancy may be due to different analysis approaches.

Samanta et al. (2021) investigated changes in BOLD response, but not functional connectivity, for the two networks separately. Their functional connectivity analysis was built on a seed-based approach which tested the interactions between a pre-defined hub of the executive control network and the rest of the brain. In contrast, we used independent component analysis to define our networks without a priori assumptions and assessed functional connectivity changes only within each of these networks. These methodological differences may have contributed to the different results of the two analyses.

The executive control network is thought to allocate attention between visuospatial tasks and is involved in goal-directed behaviors and navigation. Specifically, the posterior parietal cortex serves as a cortical integration site for hippocampally generated allocentric spatial information and egocentric spatial orientation to guide goal-directed navigation (Nitz, 2012; Whitlock et al., 2008). Additionally, there is evidence on the sequential replay or patterned reactivation of brain regions at post-task rest involved in encoding a prior experience (Hoffman & McNaughton, 2002; Staresina et al., 2013). Taken together, the evidence on the central role of the parietal cortex in spatial navigation and replay at post-task rest in task-relevant brain regions aligns with our findings. We observed increased functional connectivity in the right executive control network centered on the inferior posterior parietal lobe and sulcus at the resting state from pre- to post-task performance.

Initially, the default mode network was defined as a task-negative network that decreases in functional connectivity during task performance compared to the resting state (Buckner et al., 2008). However, using graph theory measures, Lin et al. (2017) provided a temporally resolved view on the default mode network dynamics during pre-task rest, task performance, and post-task rest. They found a decrease in efficiency, a measure of functional integration, in the default mode network at post-task rest compared to pre-task rest. Global efficiency decreased in the whole network, and local efficiency decreased specifically in the posterior cingulate cortex. Our results also demonstrate that the default mode network, with the considerable influence of the posterior cingulate cortex, decreases in functional connectivity from pre- to post-task rest. It should be pointed out that, as a result of the independent component analysis (ICA) and the template-matching procedure, the posterior cingulate cortex, the prefrontal cortex, and the lateral posterior parietal cortex contributed most strongly to the default mode network connectivity (see Supplementary Movie M2). Previous studies, including Samanta et al. (2021), also considered the hippocampus and the retrosplenial cortex functional hubs in this network (Greicius et al., 2003; Vincent et al., 2006). Together, all these regions have been proposed to form a memory network in which information is relayed from the hippocampus via the prefrontal and retrosplenial cortex to downstream parietal regions during consolidation (Genzel, 2020). However, in our study, the hippocampus and retrosplenial cortex were primarily part of a separate hippocampal network. Thus, several findings in our study that have previously been assigned to the default mode network are here discussed separately for the hippocampal network, including the hippocampus, temporal lobe regions, and the retrosplenial cortex.

Our fMRI processing pipeline was based on an ICA that created maximally independent spatial networks. Thus, we were not able to test interactions between the functional resting-state networks. Nevertheless, the opposite trends in the right executive control and default mode networks’ functional connectivity hint in the same direction as previous research. It has been shown that opposite shifts between the default mode network and the executive control network facilitate the transfer between resting and focusing attention (Goulden et al., 2014; Menon & Uddin, 2010). This transfer involves the re-allocation of resources within the brain to support stimulus-related cognitive processing. Based on our observations, the antagonism in functional connectivity between the two networks may not only be present from rest to task transitions but persists even at post-task rest and could thus reflect offline coordination of relevant areas to strengthen the encoded experience.

### The Role of the Hippocampal Network in Memory Consolidation

Through the SVD, we were able to identify two functionally relevant subnetworks within the hippocampal network, namely a primary hippocampal subnetwork centered on the hippocampus and parahippocampal cortex and a second hippocampal subnetwork primarily involving the retrosplenial cortex. These subnetworks showed distinct functional connectivity changes across the four resting-state fMRI sessions.

The primary hippocampal subnetwork increased in functional connectivity after task performance in the learning and retrieval sessions. This finding may underlie a similar resting-state consolidation as described for the executive control network. The hippocampus as initial memory storage site, has often been shown to produce reactivations that led to better memory performance after rest (Dupret et al., 2010). We also observed an increase in functional connectivity after the 90 min break for this subnetwork. At first glance, this increase across the consolidation period seems counterintuitive to the proposed mechanisms of systems consolidation (Squire et al., 2015). One would instead expect decreased involvement of the hippocampus after a consolidation period. However, one should keep in mind that the current measure is not activity per se but instead within-network connectivity.

The second hippocampal subnetwork primarily involved the retrosplenial cortex but also medial temporal regions. In the literature, the hippocampus, temporal lobe regions, and the retrosplenial cortex are often regarded as part of the default mode network because they show strong coherence during rest (Greicius et al., 2003; Vincent et al., 2006). However, recently a ‘medial temporal’ subnetwork has been identified, whose decrease in connectivity over sleep was associated with increased spatial memory performance (Barnett et al., 2020; van Buuren et al., 2019). Core regions of this network include the medial temporal lobe and the retrosplenial cortex. These structures correspond to our second hippocampal subnetwork. Consequently, our second hippocampal subnetwork may play a unique role in memory consolidation (de Sousa et al., 2019; van Buuren et al., 2019).

Functionally, the second hippocampal subnetwork decreased in connectivity across the four resting-state sessions but only for participants in the sleep condition. The medial temporal lobe and retrosplenial cortex have been termed relevant for spatial navigation in humans, and rodent models (Epstein et al., 2017; Peigneux et al., 2004) and are functionally modulated by sleep, with a gradual decoupling during deep sleep (Spoormaker et al., 2010). Thus, the second hippocampal subnetwork may be coordinating the brain-wide consolidation mechanisms involving spindle-slow oscillation activity during sleep (Cowan et al., 2020; Navarro-Lobato & Genzel, 2020). Indeed, we show that our second hippocampal subnetwork centered on the retrosplenial cortex is significantly modulated by sleep and positively relates to spindle-slow oscillation coupling power, which also predicted memory performance. The coupling of ripples, spindles, and slow oscillations may be a mechanism by which memories are consolidated during sleep from the initial hippocampal storage to downstream areas, such as the posterior parietal cortex – via the retrosplenial cortex (Genzel, 2020; Hennies et al., 2016). Our second hippocampal subnetwork may be the neural substrate through which sleep exerts its beneficial effects on memory consolidation, in addition to a general form of consolidation in the primary hippocampal subnetwork discussed earlier. In summary, we show that functional connectivity patterns in the brain underlying the consolidation of memories may not simply be confined to the established resting-state networks. Using a data-driven approach, we identified variance-based subnetworks. Specifically, we differentiated the hippocampal network into a hippocampus-centered sleep-independent component and a retrosplenial-centered sleep-dependent component. Both of these networks may be involved in spatial memory consolidation. Nonetheless, only the second hippocampal subnetwork may underlie the beneficial effects of sleep on memory formation, as is shown by its association with spindle-slow oscillation coupling power that predicted memory performance. Our findings provide a more refined view on the hippocampal network in memory consolidation, in which its resting-state functional connectivity differentially changes across learning, consolidation, and retrieval of a spatial memory task in a one-day paradigm.

### The Role of Precise Spindle-Slow Oscillation Coupling in Memory Consolidation

In the framework of systems memory consolidation, the interplay between sharp wave-ripples, spindles, and slow oscillations has been identified as a pivotal mechanism to regulate the communication between the hippocampus and task-relevant neocortical regions during sleep. Sleep oscillation coupling may gradually redistribute the connectivity between these areas and strengthen the encoded experience in extra-hippocampal, predominantly neocortical networks (Genzel, 2020; Genzel, Kroes, et al., 2014). In line with previous findings, we highlight that centro-parietal fast spindles preferentially occur during the slow oscillation’s up-state (i.e., depolarized state) and frontal slow spindles synchronize mostly in the transition to the slow oscillation’s down-state (i.e., hyperpolarized state; (Muehlroth et al., 2019)). The exact timing and consistency of the triple coupling between ripples, spindles, and slow oscillations has been suggested to be crucial for successful memory consolidation (Maingret et al., 2016). Therefore, spindles coinciding with hippocampal sharp wave-ripples during the slow oscillation up-state might represent a mechanism that facilitates the transfer of memory-related information from the hippocampus to the neocortex (Siapas & Wilson, 1998). Indeed, among others (Barakat et al., 2011), our results suggest that spindle-slow oscillation coupling in humans is positively associated with memory performance. Specifically, higher spindle-slow oscillation coupling power was related to better memory performance.

It has also been suggested that spindle slow-oscillation coupling may be more pronounced when sleep immediately follows a learning experience (Schmidt et al., 2006). However, we did not observe a difference in spindle-slow oscillation coupling between the sleep test session immediately after learning and the sleep control session two weeks later with no explicit learning task beforehand. There are several reasons for the lack of differences between the sleep test and control sessions. First, in previous studies participants slept a whole night. In this period, more data can be acquired compared to our relatively short nap of 90 min, consisting of roughly one sleep cycle. The amount of data may have increased the statistical power to detect even small effects in spindle-oscillation coupling related to learning. Next, we did not control for unspecific effects of learning on the day of the sleep control session. Thus, subjects possibly consolidated other learning experiences during the nap. These experiences could have evoked similar patterns of spindle-slow oscillation coupling during the sleep control and test sessions. Finally, and most probably, our sequence of the two sleep sessions was not counterbalanced. This was initially done to keep subjects blinded to the experimental condition at the test session. However, this also led to participants being more familiar with the sleep laboratory at the sleep control session when they encountered the setting a second time. Therefore, participants may have slept longer and better, as is indicated by higher total sleep time and fast spindle amplitude in the sleep control session.

In summary, we observed a temporally specific association between spindles and slow oscillations. Moreover, higher spindle-slow oscillation coupling power was related to better memory performance as well as decrease in functional connectivity in the second hippocampal network. Even though the coupling did not differ between sleep test and control sessions, the relation between spindle-slow oscillation coupling power and memory performance was only established for the sleep test session but not the sleep control session. Therefore, in the following, we will interpret our sleep-EEG results and their relation to the resting-state fMRI results with respect to the spatial memory task at the test day.

### Brain-Wide Offline Consolidation across Memory Conditions

The traditional view on spatial memory consolidation divides memories into hippocampus-dependent and ‘non-hippocampus-dependent’ subtypes. Accordingly, allocentric spatial memories have been proposed to depend on the hippocampus and their consolidation may profit from sleep. However, egocentric spatial memories rely on the striatum and should therefore be less sleep-dependent. Previous research provided evidence for this dissociation (Albouy et al., 2015; Hagewoud et al., 2010). There was no significant interaction between sleep and allocentric versus egocentric groups, and we also found no differences in the sleep-EEG or resting-state fMRI results. In the analysis of our dataset during task performance, Samanta et al. (2021) found a change in activity across multiple brain regions after sleep in humans and rats. Interestingly, this change was also the same for allocentric and egocentric training conditions. Irrespective of the learning strategy, task-related activity increased in areas of the executive control network and decreased in the default mode network (incl. the hippocampus) over sleep in humans. These findings argue against a strict separation into hippocampus-dependent and ‘non-hippocampus-dependent’ memories.

Recent evidence supports the involvement of the hippocampus in non-hippocampus-dependent memories. A rodent study tested the effects of sleep on novel-object recognition memory. The encoding and retrieval of these tasks essentially relied on the perirhinal cortex and not the hippocampus (Sawangjit et al., 2018). Importantly, sleep-specific enhancement of object-recognition memory required hippocampal activity during post-encoding sleep, but the hippocampus was not involved during remote recall. Another study could show the same in human subjects (Schapiro et al., 2019). Furthermore, Ekstrom et al. (2017) outlined a network model of human spatial navigation in which several largely overlapping brain regions encode allocentric and egocentric information. Considering this network perspective, resulting memory representations of the two reference frames seem to be intertwined as well. It has been shown that the ventral striatum integrates inputs from the hippocampus, prefrontal cortex, and related subcortical structures to perform goal-directed behavior. Later memory reactivations in the striatum were temporally close to hippocampal ripples (Pennartz et al., 2011; Pennartz et al., 2004). Together, these findings suggest that interactions in allocentric and egocentric memory networks are more complex than originally thought. Perhaps, consolidation mechanisms at sleep and rest are even independent of the learning strategy. This domain generality in memory consolidation could enable flexibility and adaptability for information access, use, and reconsolidation.

## Conclusion

This study aimed to detect representations of spatial memory consolidation at different brain states (sleep and rest). We quantified memory consolidation mechanisms during sleep using spindle-slow oscillation coupling and observed changes in resting-state functional connectivity across learning, consolidation, and retrieval in the executive control network, the default mode network, and the hippocampal network. A data-driven analysis allowed us to further divide the hippocampal network into a sleep-dependent and a sleep-independent subnetwork. We showed that spindle-slow oscillation coupling power was associated with deceased functional connectivity in the sleep-dependent hippocampal subnetwork, which included the retrosplenial cortex, and better memory performance.

## Acknowledgments

The study was supported by a Branco Weiss Fellowship – Society in Science – awarded to L.G.

## Author contributions

L.B. performed the analysis and wrote the first draft of the article, A.S. acquired the data, F.W. D.R.P and M.D. supervised the analysis, R.S. designed the virtual watermaze task, L.G. designed the study, supervised the data acquisition and analysis and co-wrote the article. All authors contributed to corrections in the article.

## Supplementary Materials

**Figure S1.**
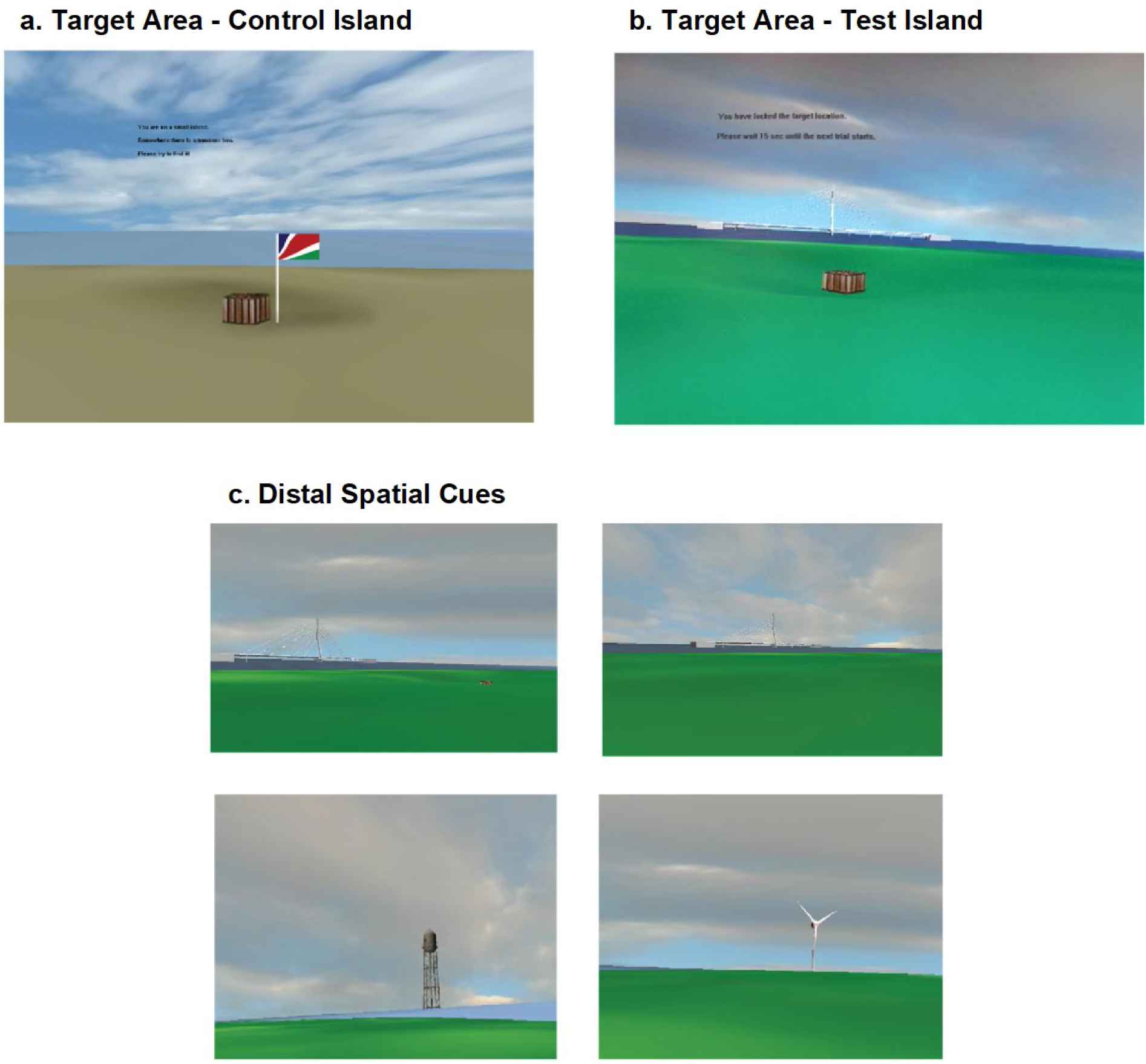
Snapshots of the virtual task environment. **a**, Representation of the cued island, a plain brown island with no surrounding distal landmarks. The cue is a flag that is visible from a distance and keeps changing its position each trial. **b**, Representation of the target quadrant of the hidden island, a green island surrounded by four landmarks (see c). The location of the box is fixed on this island and is located on an indentation in the virtual surface and only visible from close distance. **c**, Distal orientation landmarks: a bridge, a sailboat, a windmill and a light house. The upper left picture in the panel displays the target quadrant from a distance. The box is only visible when the subject is close to the target location. Reproduced with permission from Samanta et al. (2021).

**Table T1.**
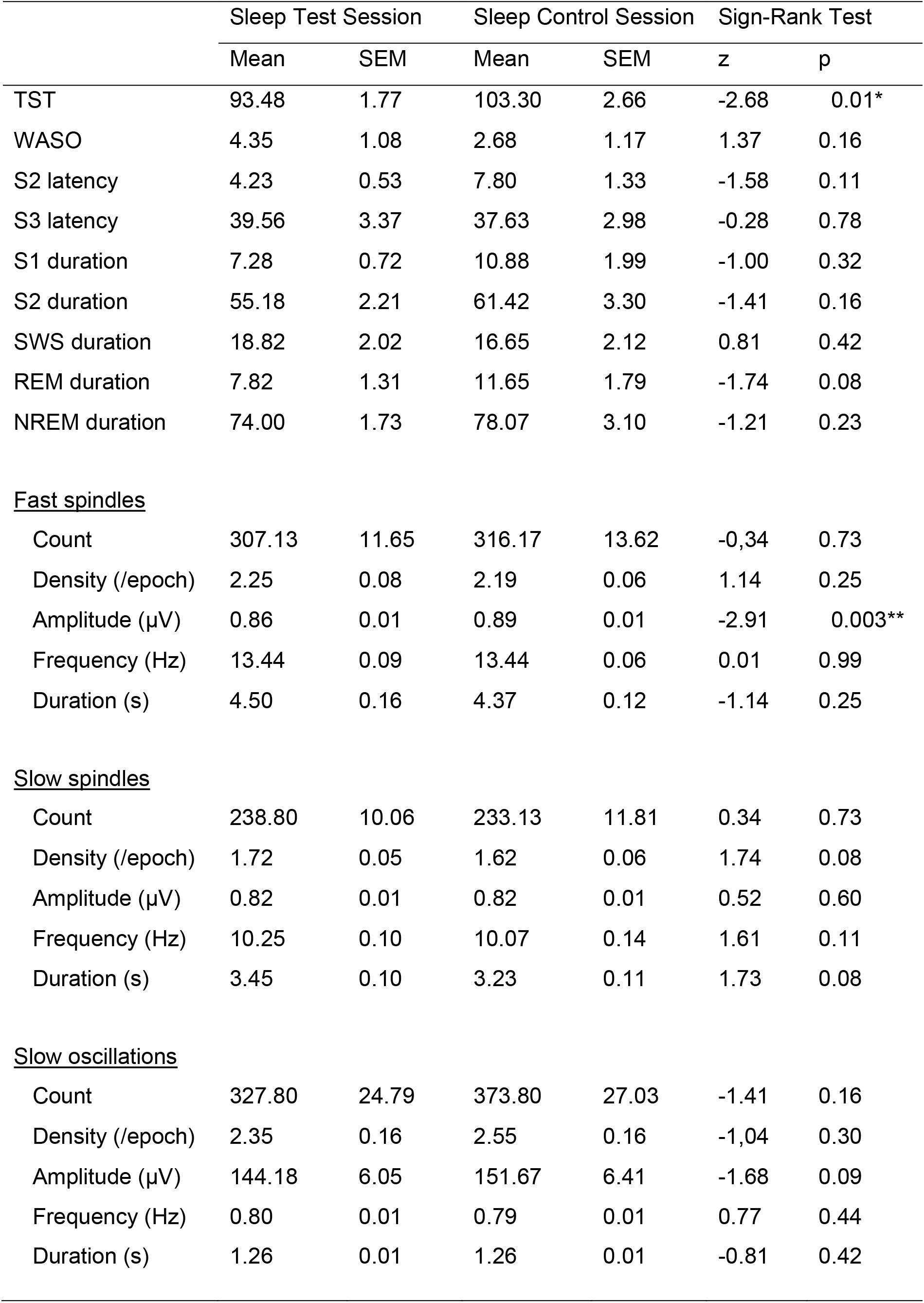
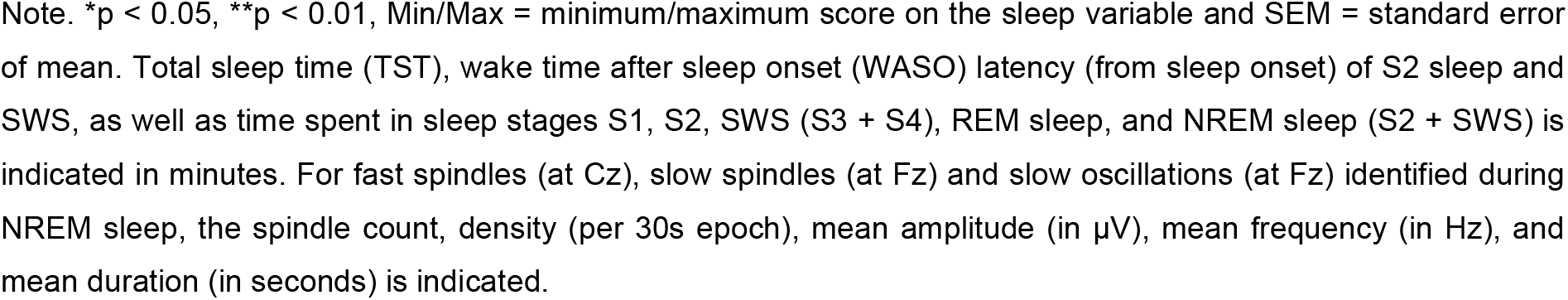
Sleep Descriptives and Oscillation Properties across Sleep Sessions

|  | Sleep Test Session |  | Sleep Control Session |  | Sign-Rank Test |  |
| --- | --- | --- | --- | --- | --- | --- |
|  | Mean | SEM | Mean | SEM | z | p |
| TST | 93.48 | 1.77 | 103.30 | 2.66 | -2.68 | 0.01* |
| WASO | 4.35 | 1.08 | 2.68 | 1.17 | 1.37 | 0.16 |
| S2 latency | 4.23 | 0.53 | 7.80 | 1.33 | -1.58 | 0.11 |
| S3 latency | 39.56 | 3.37 | 37.63 | 2.98 | -0.28 | 0.78 |
| S1 duration | 7.28 | 0.72 | 10.88 | 1.99 | -1.00 | 0.32 |
| S2 duration | 55.18 | 2.21 | 61.42 | 3.30 | -1.41 | 0.16 |
| SWS duration | 18.82 | 2.02 | 16.65 | 2.12 | 0.81 | 0.42 |
| REM duration | 7.82 | 1.31 | 11.65 | 1.79 | -1.74 | 0.08 |
| NREM duration | 74.00 | 1.73 | 78.07 | 3.10 | -1.21 | 0.23 |
| <u>Fast spindles</u> |  |  |  |  |  |  |
| Count | 307.13 | 11.65 | 316.17 | 13.62 | -0.34 | 0.73 |
| Density (/epoch) | 2.25 | 0.08 | 2.19 | 0.06 | 1.14 | 0.25 |
| Amplitude (μV) | 0.86 | 0.01 | 0.89 | 0.01 | -2.91 | 0.003** |
| Frequency (Hz) | 13.44 | 0.09 | 13.44 | 0.06 | 0.01 | 0.99 |
| Duration (s) | 4.50 | 0.16 | 4.37 | 0.12 | -1.14 | 0.25 |
| <u>Slow spindles</u> |  |  |  |  |  |  |
| Count | 238.80 | 10.06 | 233.13 | 11.81 | 0.34 | 0.73 |
| Density (/epoch) | 1.72 | 0.05 | 1.62 | 0.06 | 1.74 | 0.08 |
| Amplitude (μV) | 0.82 | 0.01 | 0.82 | 0.01 | 0.52 | 0.60 |
| Frequency (Hz) | 10.25 | 0.10 | 10.07 | 0.14 | 1.61 | 0.11 |
| Duration (s) | 3.45 | 0.10 | 3.23 | 0.11 | 1.73 | 0.08 |
| <u>Slow oscillations</u> |  |  |  |  |  |  |
| Count | 327.80 | 24.79 | 373.80 | 27.03 | -1.41 | 0.16 |
| Density (/epoch) | 2.35 | 0.16 | 2.55 | 0.16 | -1.04 | 0.30 |
| Amplitude (μV) | 144.18 | 6.05 | 151.67 | 6.41 | -1.68 | 0.09 |
| Frequency (Hz) | 0.80 | 0.01 | 0.79 | 0.01 | 0.77 | 0.44 |
| Duration (s) | 1.26 | 0.01 | 1.26 | 0.01 | -0.81 | 0.42 |

Note. \* $p < 0.05$ , \*\* $p < 0.01$ , Min/Max = minimum/maximum score on the sleep variable and SEM = standard error of mean. Total sleep time (TST), wake time after sleep onset (WASO) latency (from sleep onset) of S2 sleep and SWS, as well as time spent in sleep stages S1, S2, SWS (S3 + S4), REM sleep, and NREM sleep (S2 + SWS) is indicated in minutes. For fast spindles (at Cz), slow spindles (at Fz) and slow oscillations (at Fz) identified during NREM sleep, the spindle count, density (per 30s epoch), mean amplitude (in $\mu\text{V}$ ), mean frequency (in Hz), and mean duration (in seconds) is indicated.

**Figure S2.**
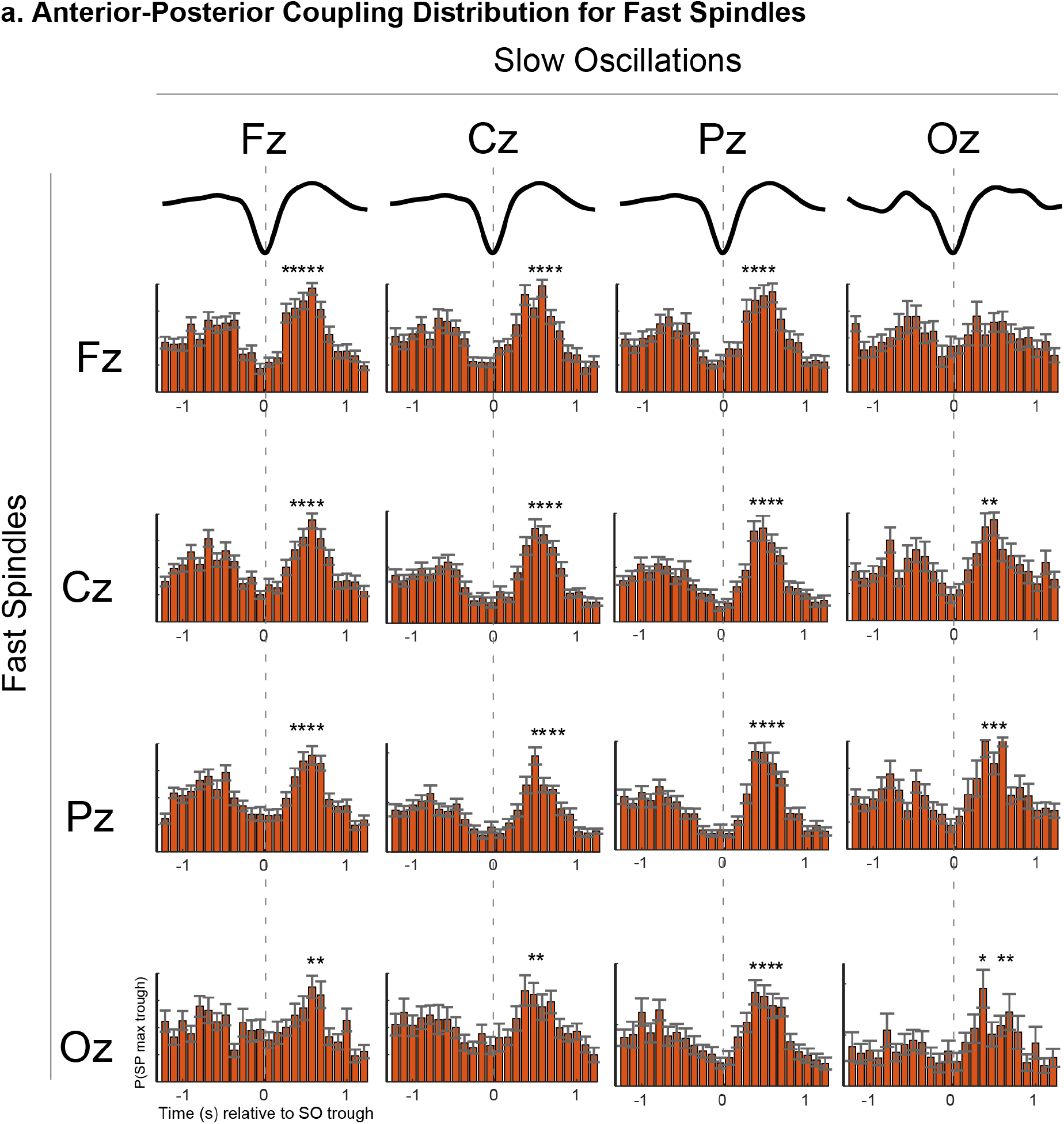
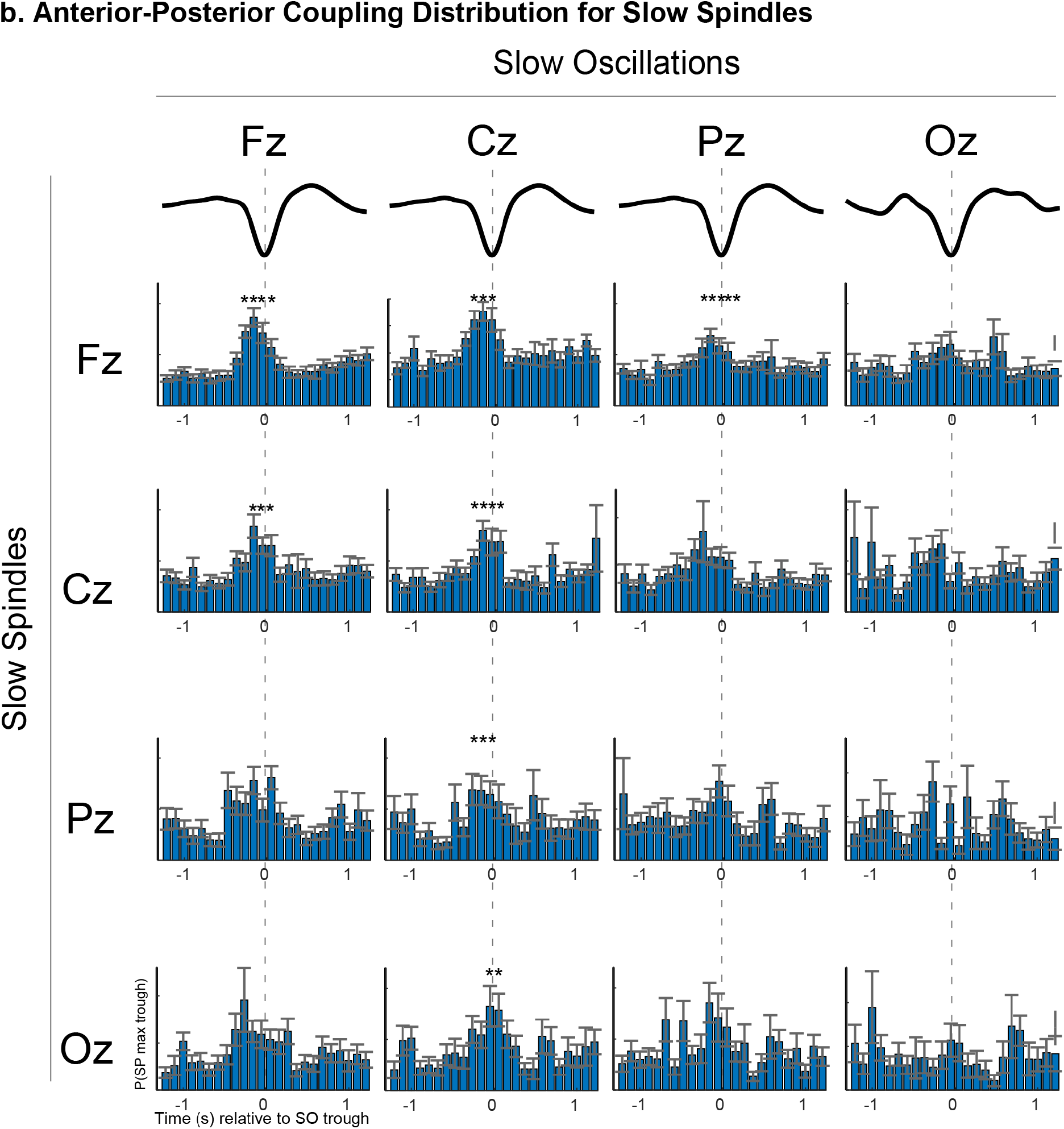
Central anterior-posterior distribution of the temporal spindle-slow oscillation coordination at sleep test session. **a,** Fast spindle-slow oscillation PETHs as described in in Fig. 2b plotted for fast spindles (rows) and slow oscillations (columns) for all possible combinations of midline channels. Asterisks indicate significantly increased spindle occurrence contrasted with the reference distribution (cluster-based permutation test, cluster α < 0.05, positive clusters only). Vertical dashed lines mark the slow oscillation trough. Average slow oscillations are shown for each channel at the top of the plot. Error bars of 100-ms time bins = SEM. **b**, Same as **a** but for slow spindles. Note that the coordination between spindles and slow oscillations at this temporal resolution does depend negligibly on the combination of channel selection for channels Fz and Cz. Generally, fast spindles occur mostly during the slow oscillation up-state and slow spindles mostly in the transition to the slow oscillation down-state. The pattern of fast spindle-slow oscillation coupling is less pronounced for more posterior channels and for distant pairs of channels. Slow spindle-slow oscillation coupling is rather confined to frontal channels.

**Figure S3.**
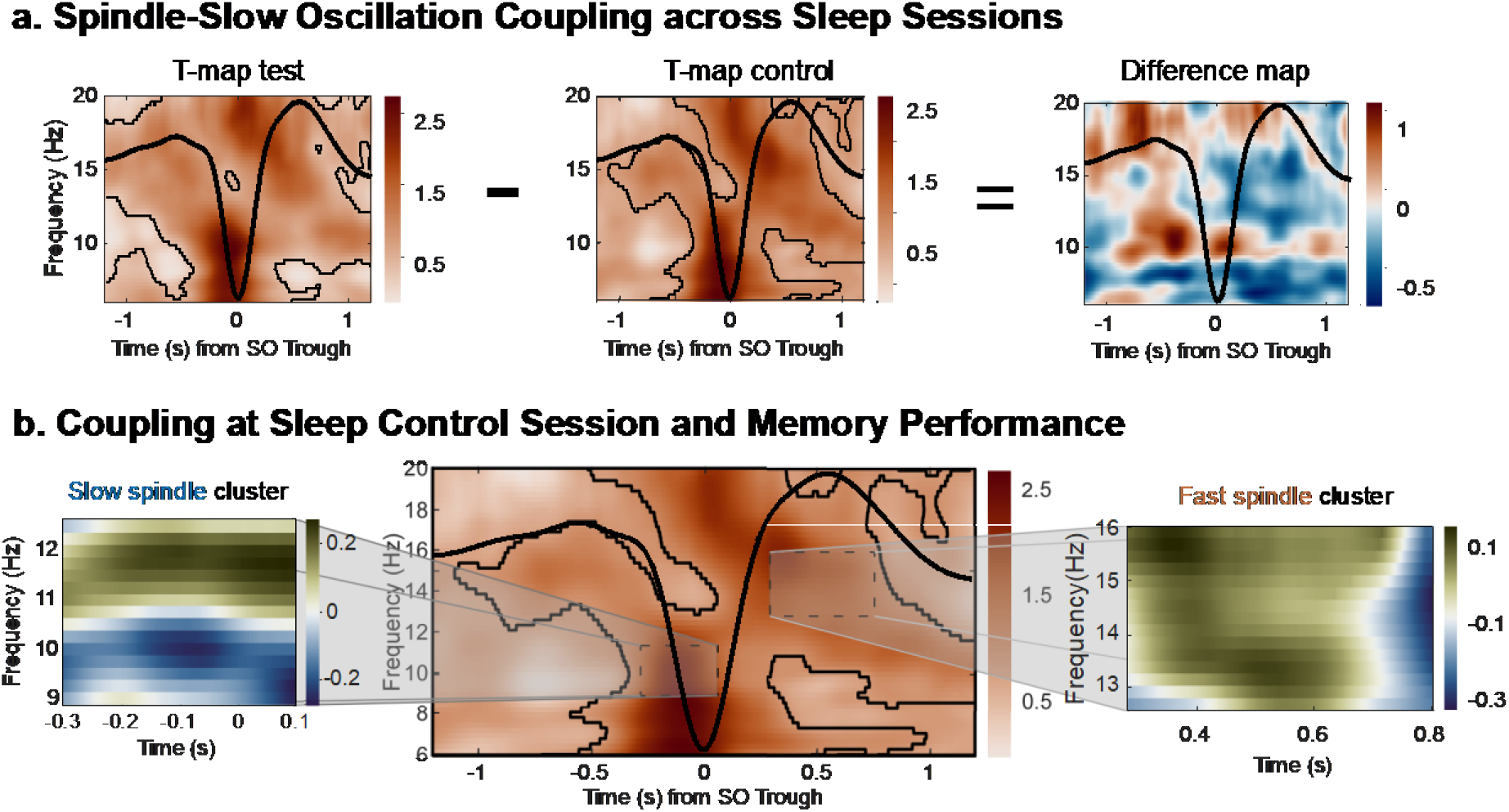
Fast spindle-slow oscillation coupling power in the sleep control session is not associated with memory performance. **a**, Power modulations in across the sleep test session and sleep control session. Differences in superlet power for slow oscillation trials (respective trough ± 1.2 s) compared to baseline trials without slow oscillations are depicted (in t-score units) for the two sleep sessions separately. T-map of the control session is subtracted from that of the test session to obtain the difference map. The average frontal slow oscillation for each session is shown in black. In both sessions, EEG activity is modulated as a function of the slow oscillation phase but there is no difference between the two sessions. Significant clusters are outlined in black (cluster-based permutation test, cluster α < 0.05) **b**, Correlation of spindle-slow oscillation power modulations and memory performance. *left:* T-map as outlined in Figure 4c but for the sleep control session including subjects with spindle-slow oscillation coupling above chance level at sleep test session (see Fig. 4b; slow spindles and fast spindle reference windows for later analyses highlighted by dashed black line). *right:* Fast spindle window with correlation values between slow oscillation-specific EEG activity and memory performance obtained by contrasting the correlation. Note that in the sleep control session there is neither the fast spindle cluster nor the slow spindle cluster is significantly correlated with memory performance at test day. This is different to the two spindle cluster of the sleep test session (see Fig 4c).

**Figure S4.**
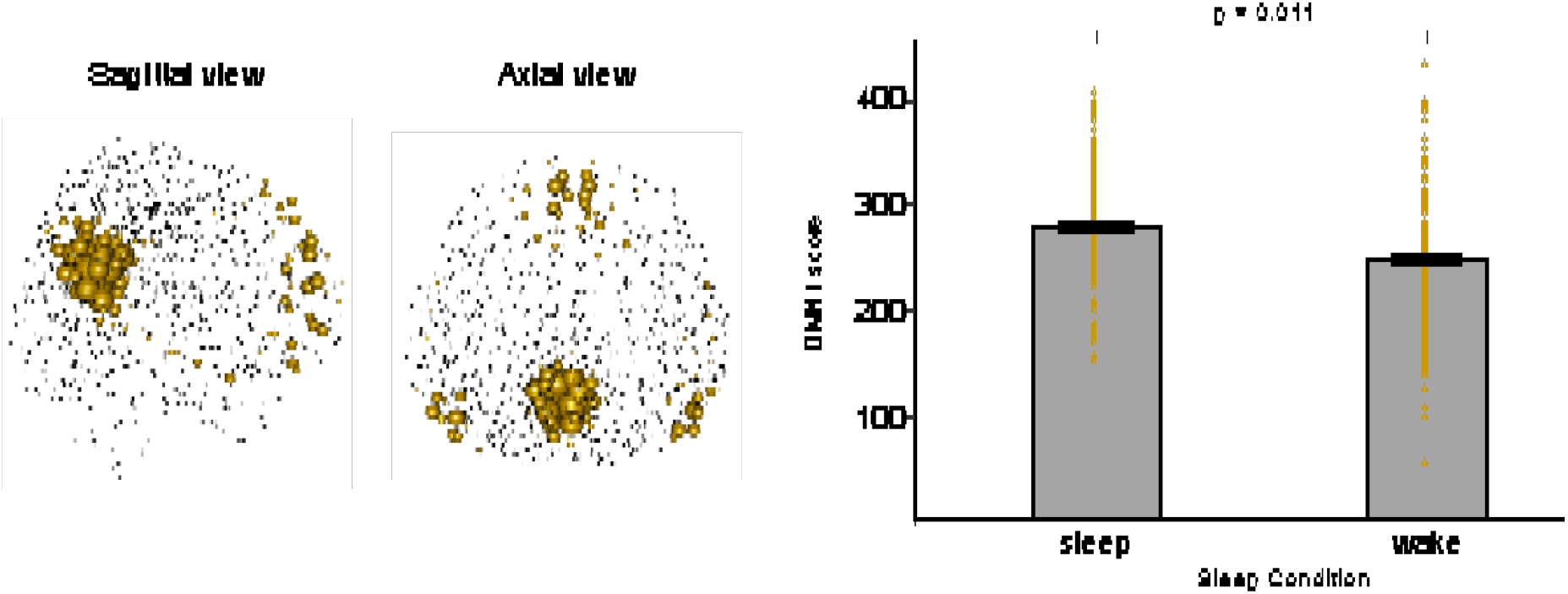
Difference between sleep conditions in the primary mode of the default mode network. *left:* Sagittal and axial representation of the primary mode in the default mode network as shown in Fig. 5. *right:* Partial results of a linear mixed-effects model with session as within-subject factor, sleep and memory as between-subjects factor and subject*session as random factor.

